# Epitranscriptomic reprogramming is required to prevent stress and damage from acetaminophen

**DOI:** 10.1101/2021.08.16.456530

**Authors:** Sara Evke, J. Andres Melendez, Qishan Lin, Thomas J. Begley

## Abstract

Epitranscriptomic marks, in the form of enzyme catalyzed RNA modifications, play important gene regulatory roles in response to environmental and physiological conditions. However, little is known with respect to how pharmaceuticals influence the epitranscriptome. Here we define how acetaminophen (APAP) induces epitranscriptomic reprogramming and how the writer Alkylation Repair Homolog 8 (Alkbh8) plays a key gene regulatory role in the response. Alkbh8 modifies tRNA selenocysteine (tRNA^Sec^) to translationally regulate the production of glutathione peroxidases (Gpx’s) and other selenoproteins, with Gpx enzymes known to play protective roles during APAP toxicity. We demonstrate that APAP increases toxicity and markers of damage, and decreases selenoprotein levels in *Alkbh8* deficient mouse livers, when compared to wildtype. APAP also promotes large scale reprogramming of 31 RNA marks comprising the liver tRNA epitranscriptome including: 5-methoxycarbonylmethyluridine (mcm^5^U), isopentenyladenosine (i^6^A), pseudouridine (Ψ), and 1-methyladenosine (m^1^A) modifications linked to tRNA^Sec^ and many others. Alkbh8 deficiency also leads to wide-spread epitranscriptomic dysregulation in response to APAP, demonstrating that a single writer defect can promote downstream changes to a large spectrum of RNA modifications. Our study highlights the importance of RNA modifications and translational responses to APAP, identifies writers as key modulators of stress responses *in vivo* and supports the idea that the epitranscriptome may play important roles in responses to pharmaceuticals.

## Introduction

Epitranscriptomic marks catalyze RNA modifications and are important gene regulatory signals, with defects linked to disrupted gene expression, disease onset and disease progression (Kadumuri and Janga 2018). Modifications on mRNA, rRNA and tRNA have been shown to be dynamically regulated to allow cells to survive different stressors and adapt to changes in cellular physiology (Cai et al. 2016; Chan et al. 2012; Endres, Dedon, and Begley 2015; Gu, Begley, and Dedon 2014). In addition, the location and levels of specific RNA modifications have been shown to drive gene expression. tRNAs are the most heavily modified RNA species, and RNA modifications are critical for regulating tRNA structure, function and stability. Writer enzymes can catalyze or add tRNA modifications, while eraser enzymes can remove modifications and reader enzymes bind and recognize modifications. Chemical modifications on tRNAs provide regulation of structure and function, while those occurring in the anticodon loop positions 34 to 37 can regulate translation and fidelity. The wobble position (34) in the anticodon stem loop of tRNA allows pairing to occur with more than one nucleoside, allowing for a single tRNA to decode multiple codons with different 3’ nucleosides (Agris, Vendeix, and Graham 2007). Wobble position modifications to uridine (U), cytidine (C), guanosine (G) and adenosine (A) have all been reported and can include simple methylations to more elaborate chemical additions, which can regulate anticodon-codon interactions while preventing translational errors (Agris et al. 2017, 2018; Agris, Vendeix, and Graham 2007; Novoa, Pavon-eternod, and Pan 2012).

While most mammalian tRNAs have 3 to 17 positions modified (Pan 2018), the tRNA for selenocysteine (tRNA^Sec^) is unique because it only has 4 positions modified. tRNA^Sec^ modifications and writers include the 5-methoxycarbonylmethyluridine (mcm^5^U) and the ribose-methylated derivative, 5-methoxycarbonylmethyl-2’-O-methyluridine (mcm^5^Um) at position U34 which are dependent on Alkylation repair homolog 8 (Alkbh8) along with its accessory protein tRNA methyltransferase 112 (Trm112). The isopentenyladenosine (i^6^A) modification at position A37 is catalyzed by tRNA isopentenyltransferase 1 (Trit1) (Fradejas et al. 2013), while the pseudouridine (Ψ) modification at position U55 is catalyzed by pseudouridine synthase 4 (Pus4) also known as TruB pseudouridine synthase family member 1 (Trub1) (Roovers et al. 2006), and the 1-methyladenosine (m^1^A) modification at position A58 is catalyzed by the catalytic subunit of tRNA (adenine-N_1_-)-methyltransferase (Trmt61A) and RNA binding protein non-catalytic subunit tRNA (adenine (58)-N(1))-methyltransferase (Trm6) (M. Wang et al. 2016). Alkbh8, mcm^5^U and mcm^5^Um have been shown to play a critical roles in the translation of selenoproteins (Born et al. 2011; Van Den Born et al. 2011; D. Fu et al. 2010; Pastore et al. 2012; Songe-Moller et al. 2010). Selenoprotein synthesis is orchestrated by interactions between cis RNA elements and trans proteins and specifically modified tRNA to recode a UGA stop codon. Thus, the UGA selenocysteine (Sec) codon found within selenoprotein genes works with Alkbh8-modified tRNA, as well as other factors, to promote the translation of selenoproteins. There are 25 selenoproteins in human systems with rodents having only 24, with glutathione peroxidase 6 (Gpx6) being the discrepant selenoprotein (Regina et al. 2016). Some examples of selenoproteins include glutathione peroxidase (Gpx)1-4, thioredoxin reductases (TrxR) 1-3, selenoprotein S (SelS), selenoprotein K (SelK), and selenoprotein P (SelP).

Selenoproteins play roles in stress responses, regulating metabolism, immunity, and in embryonic vitality and development (Anouar et al. 2018; Hatfield et al. 2006; Hofstee, Cuffe, and Perkins 2020; Pappas, Zoidis, and Chadio 2019; Pitts et al. 2014; Rundlöf and Arner 2004; Shrimali et al. 2008). Many selenoproteins serve as antioxidant enzymes to mitigate damage caused by reactive oxygen species (ROS) (Arbogast and Ferreiro 2010; Benhar 2018; Chung et al. 2009; Couto, Wood, and Barber 2016; Eckers et al. 2013; Huang et al. 2020; Leonardi et al. 2019; Steinbrenner and Sies 2009; Zoidis et al. 2018). A GeneTrap mouse deficient in *Alkbh8* (*Alkbh8^Def^*) has been used to generate embryonic fibroblasts (MEFs) and these cells have been used to demonstrate that writer defects promote increased levels of intracellular ROS and DNA damage and reduced selenoprotein expression. *Alkbh8^Def^* MEFS also display a proliferative defect, accelerate senescence and fail to increase selenoprotein levels in response hydrogen peroxide (H_2_O_2_) (Endres et al. 2015a; Lee et al. 2020). *Alkbh8^Def^* mice also have increased lung inflammation and a disruption in lung glutathione levels (Leonardi et al. 2020). Pulmonary exposure of *Alkbh8^Def^* mice to naphthalene, the common polyaromatic hydrocarbon found in mothballs, increased stress markers and lung damage, when compared to their wildtype (WT) counterparts (Leonardi et al. 2020). Overall, these findings demonstrate that the epitranscriptomic writer *Alkbh8* plays a vital role in protection from environmental stressors.

Little is known with respect to how the epitranscriptome responds to pharmaceutical stress. Acetaminophen (APAP) is one of the most prevalent over the counter drugs due to its analgesic and antipyretic properties and is consumed regularly by over 60 million Americans each week. APAP is a safe and effective drug that combats various ailments. However, in the United States nearly 50% of all cases of acute liver failure are due to an excessive use of APAP (Yoon et al. 2016). Gpx proteins have been linked to preventing APAP toxicity by providing protection from its metabolized forms (Mirochnitchenko et al. 1999). The majority of APAP (85-90%) is acted upon by a phase II metabolism where APAP is chemically altered by UDP-glucuronosyl transferases (Ugt) and sulfotransferases (Sult) and then converted to glucouronidated and sulfated metabolites that are eliminated in urine. A small amount (∼2%) of APAP is excreted in urine without modification and the remaining is biometabolized by the cytochrome enzyme, Cyp2e1, which results in the highly reactive, toxic metabolite *N*-acetyl-para-benzoquinone imine (NAPQI) (Mcgill and Jaeschke 2014). Sufficient glutathione (GSH) in hepatocytes will reduce the reactive NAPQI and promote excretion in the urine. However excessive amounts of NAPQI can increase in oxidative stress, decrease GSH, and promote mitochondrial dysfunction leading to depletion of adenosine triphosphate (ATP) stores (Clark et al. 2012).

Selenoproteins have been shown to provide protection from APAP toxicity. Mirochnitchenko et al. demonstrated that glutathione peroxidase injection provided complete protection in mice administered an acute toxic dose of APAP (Mirochnitchenko et al. 1999). GPX3 overexpressing cells are more resistant to APAP toxicity than cells whose GPX3 levels were decreased by siGpx3 (Kanno et al. 2017). Human GPX and mouse Gpx proteins are clearly linked to preventing APAP toxicity. As selenoprotein synthesis and activity are reliant on RNA modifications that drive selenocysteine utilization, we hypothesized that Alkbh8-regulated epitranscriptomic marks would be involved in the response to APAP. Here we test the hypothesis that the epitranscriptomic writer, Alkbh8 protects against APAP toxicity by catalyzing tRNA modifications that regulate the translation of stress response genes. We have phenotypically characterized liver tissue from both C57BL/6 WT and *Alkbh8^Def^* treated mice with either saline (vehicle control) or toxic (600 mg/kg) APAP under both acute (6h) and sub-chronic (4 daily injections) exposure conditions. When challenged with APAP, WT liver tissue showed an increase in many selenoproteins whereas the *Alkbh8^Def^* liver tissues were disrupted in their expression and show classic hall marks of stress and damage, including increased expression of the oxidative stress marker, 8-isoprostane, and liver damage marker, alanine transaminase (ALT). Transcriptional responses to acute APAP were similar between the genotypes; however, protein level differences were detected, which is indicative of post-transcriptional changes reflective of epitranscriptomic regulation. After a single dose of APAP, we have discovered that there is significant change in the levels of 31 tRNA modifications in response to APAP. Increases in the levels of tRNA modifications specific to tRNA^Sec^ and many other tRNA isoacceptors was identified, with dysregulation of the many APAP dependent marks observed in *Alkbh8^Def^* mice. Our study is one of the first to characterize the roles of epitranscriptomic marks and writers in response to APAP, and supports the idea that the epitranscriptome adapts to and modifies the effects of pharmaceuticals.

## Material and Methods

### 2.1. Animal Experiments

Animal studies were carried out in strict accordance with recommendations in the guide for the Care and Use of Laboratory Animals of the National Institutes of Health. The protocol was approved by the University at Albany Institutional Animal Care and Use Committee (Albany, NY), protocol #17-016/#20-014 for breeding and #18-010 for APAP exposure. Two homozygous strains of mice on the C57BL/6J background were bred, WT and *Alkbh8^Def^*. Wild-type mice were purchased from Taconic Biosciences (Rensselaer, NY). *Alkbh8^Def^* were produced through an insertional mutagenesis approach targeting the *Alkbh8* gene in the parental E14Tg2a.4 129P2 embryonic stem cell line, as previously described (Endres et al., 2015). *Alkbh8^Def^* mice were produced using a vector containing a splice acceptor sequence upstream of a β (β-galactosidase/neomycin phosphotransferase fusion), which was inserted into intron 7 at chromosome position 9:3349468-3359589, creating a fusion transcript of Alkbh8-β-geo (Stryke et al. 2003). Multiplex qRT-PCR and relative cycle threshold analysis (ΔΔCT) on genomic DNA derived from tail biopsies were used to determine animal zygosity for *Alkbh8^Def^*, using TaqMan oligonucleosides specific for neomycin (Neo, target allele) and T-cell receptor delta (Tcrd, endogenous control). Neo forward (5’-CCA TTC GAC CAC CAA GCG-3’), Neo reverse p (5’-AAG ACC GGC TTC CAT CCG-3’), Neo Probe (5’-FAM AAC ATC GCA TCG AGC GAG CAC GT TAMRA-3’), Tcrd forward (5’-CAG ACT GGT TAT CTG CAA AGC AA-3’), Tcrd reverse (5’-TCT ATG CAA GTT CCA AAA AAC ATC-3’), and Tcrd probe (5’-VIC ATT ATA ACG TGC TCC TGG GAC ACC C TAMRA −3’) primers were used for genotyping.

For all animal studies, male mice between 8-12 weeks of age were used. Females were not used, as they have been shown to be less susceptible to APAP liver injury due to their accelerated recovery of hepatic glutathione (GSH) levels (Du et al. 2014). Mice were sacrificed via CO_2_ asphyxiation and in some cases, followed by non-survival vena cava puncture and necropsy of the liver. Whole blood was allowed to clot for 30 minutes at 25°C and organs were flash-frozen and placed in −80°C for storage.

### 2.2. APAP solution preparation and intraperitoneal injection (I.P.)

APAP and Dulbecco’s Phosphate Buffered Saline (PBS) were purchased from Sigma-Aldrich (St. Louis, MO). A 5 mg/mL (w/v) solution of APAP was made in a 15 mL conical tube with 10 mL of PBS and 0.5 g of APAP, due to lower solubility property of APAP, the solution was boiled until dissolved and then cooled to room temperature before injection. Solutions were made fresh each day. For intraperitoneal injections (I.P.) mice were first weighed and then placed under 5% isoflurane gas for approximately three minutes. The mice were then manually restrained, the injection site was wiped with 70% ethanol, and either saline or 600 mg/kg APAP solution was injected into the intraperitoneal cavity in the lower left abdominal quadrant using Coviden™ Monoject™ 27g, ½ inch standard hypodermic needles (Fisher Scientific, Franklin, MA) and sterile 1 ml Coviden™ Monoject™ tuberculin syringes (Fisher Scientific, Franklin, MA). Injections for acute exposures occurred once and six hours after the exposure mice were sacrificed. For chronic conditions, injections were administered at the same time daily for four days, every 24 hours, and mice were sacrificed 24 hours post last injection. After injection, mice were observed up to one hour for recovery from anesthesia and to ensure mice were not experiencing any pain or distress. Mice were further monitored for signs of excess stress and weight loss, with any mouse losing more than 20% of its body weight sacrificed based on IACUC guidelines.

### 2.3 Alanine Transaminase (ALT) Assay

The Alanine Transaminase Colorimetric Activity Assay (Catalog no. 700260, Cayman Chemical, Ann Arbor, MI) was performed following manufacturer’s protocol to provide an indicator of liver damage. WT and *Alkbh8^Def^* were exposed to either saline or APAP as described above and sacrificed via CO_2_ asphyxiation. Cardiac puncture was then performed using Coviden™ Monoject™ 27g, ½ inch standard hypodermic needles (Fisher Scientific) and sterile 1 ml Coviden™ Monoject™ tuberculin syringes (Fisher Scientific) to collect a minimum volume of 200 µL of blood. Serum from each sample was prepared by allowing the blood to clot at 25°C for 30 minutes. Samples were centrifuged at 2,000 x g for 15 minutes at 4°C and the top serum layer was collected and stored at −80°C. Serum samples were processed based on Alanine Transaminase Colorimetric Activity Assay Kit (catalog no. 700260, Cayman Chemical) instructions, and absorbance was monitored at 340 nm over a period of 10 minutes at 1-minute intervals. Data analysis was performed and ALT activity was reported as U/L. Statistical significance was calculated utilizing three biological replicates for each exposure condition and genotype, with error bars denoting standard error of the mean and significance determined using Student’s unpaired t-test in GraphPad Prism version 9.0 (GraphPad, San Diego, CA).

### 2.4 8-isoprostane Assay

The 8-isoprostane assay (Catalog no. ab175819, Abcam, Cambridge, MA) was performed following manufacturer’s protocol. Mice were exposed to either saline or APAP as described above and sacrificed via CO_2_ asphyxiation. 1X Phosphate Buffered Saline (PBS) was injected into the portal vein and hepatic artery, to remove excess blood and other cellular debris from the liver. Once dissected from the mouse, the livers were washed in PBS and then stored at −80°C. Additional reagents that were used are 2N Sulfuric Acid Stop Solution (Catalog no. DY994, R&D Systems, Minneapolis, MN), Triphenylphosphine (TPP) (Catalog no. T84409-1G Sigma-Aldrich), and ethyl acetate (Catalog no. 270989, Sigma-Aldrich). 8-isoprostane was measured following manufacturer’s protocol with the adjustment for low sample volume. Livers were weighed and homogenized in a 500 µL solution containing 5mg TPP in dH_2_O and then acidified by adding 4 µL of glacial acetic acid to ensure pH was ∼ 4.0. Next, 500 µL of ethyl acetate was added to each sample, vortexed and centrifuged at 5,000 rpm for 5 minutes to separate the organic phase. The supernatant was collected and placed in a new conical tube and the previous step was performed one more time. The pooled organic phase supernatant was dried using nitrogen gas and then reconstituted in 20 µL 100% ethanol. Samples were further processed according to manufacturer’s protocol and then used for ELISA measurement and analyzed. Statistical significance was calculated utilizing three biological replicates for each exposure condition and genotype with error bars denoting standard error of the mean and significance determined using Student’s unpaired t-test in GraphPad Prism v9.

### 2.5. Protein Quantification Assays

Primary antibodies used for quantification were as follows: TrxR1 (catalog no. MAB7428, R&D Systems), TrxR2 (catalog no. ab180493, Abcam, Cambridge, MA), Gpx1 (Catalog no. AF3798, R&D Systems), Gpx3 (Catalog #AF4199, R&D Systems), Gpx4 (catalog no. 565320, Novus Biologicals, Centennial, CO), SelS (catalog no. HPA010025, Sigma) and Gapdh (catalog no. CB1001, Millipore Sigma). Liver tissue was homogenized in 1 mL RIPA Buffer (50 mM Tris-HCl, pH 7.4, with 150 mM NaCl, 1% TridonX-100, 0.5% sodium deoxycholate, and 0.1% sodium dodecyl sulfate) supplemented with an EDTA-free phosphatase inhibitor cocktail 2 (catalog no. P5726, Sigma) and phosphatase inhibitor cocktail 3 (catalog no. P0044, Sigma). Livers were then agitated periodically on ice for 30 minutes and then spun at 15,000 x g for 12 minutes at 4°C. The supernatant was then collected and protein levels were quantified via Bradford assay (BioRad, Portland, OR) according to manufacturer’s instructions at a monochromatic wavelength of 595 nm. Proteins were then normalized to 1.5 µg/mL concentration and added to Wes (ProteinSimple, San Jose, CA) protein analysis plates (catalog no.SM-W004) and quantified with Compass for Simple Western v2.1. Primary antibodies of interest were always multiplexed with endogenous Gapdh primary antibody to normalize chemiluminescence area under the curve. Statistical significance was calculated utilizing three biological replicates for each exposure condition and genotype, error bars denote standard error mean with statistical significance determined using Student’s unpaired t-test in GraphPad Prism v9.

### 2.6 Liquid Chromatography Tandem Mass Spectrometry (LC-MS/MS)

Small RNAs were first isolated from liver tissue using PureLink™ miRNA Isolation Kit (catalog no. K157001, Invitrogen, Waltham, MA). RNA concentration was then quantified at The Center for Functional Genomics/University at Albany using an Agilent Technologies 2100 Bioanalyzer System (Santa Clara, CA) and respective Bioanalyzer Small RNA reagents (catalog no. 5067-1548, Agilent Technologies). Purified small RNA samples were then digested using Nucleoside Digestion Mix (catalog no. M0649S, New England BioLabs, Beverly, MA). LC-MS/MS measurement of various post-transcriptional modifications were performed on a Waters XEVO TQ-S^TM^ (Waters, USA) electrospray triple quadrupole mass spectrometer configured with ACQUITY I class ultra-performance liquid chromatography, as previously described in (Basanta-Sanchez et al. 2016). The capillary voltage was set at 1.0 kV with extraction cone of 14 V. Nitrogen flow was maintained at 1000 l/h and desolvation temperature at 500 °C. The cone gas flow was set to 150 l/h and nebulizer pressure to 7 bars. Each individual nucleoside modification was characterized by single infusion in a positive mode ionization over an m/z range of 100-500 amu. Further nucleoside characterization was performed using Waters software part of Intellistart MS/MS method development where a ramp of collision and cone voltages is applied to find optimal collision energy parameters for the daughter ions. To quantify various RNA modified nucleosides, calibration curves were prepared for these modified nucleosides including adenosine, cytidine, guanosine and uridine. [^13^C][^15^N]-G (1 pg/µL) was used as an internal standard.

A method to extract peak areas from raw data to allow quantification was developed using a combination of instrument manufactures’ suites, MassLynx V4.1 and TargetLynx (Waters, USA). These methods allowed extraction of information to produce calibration curves from each RNA modification standard. In addition, these programs were used to extract the peak areas to be extrapolated on the standard calibration curves for quantification of RNA modifications. In house Python script combined with Originlab software suite 2017 was used to quantify various RNA modifications. Heat maps were generated using MORPHEUS versatile matrix visualization and analysis software (https://software.broadinstitute.org/morpheus). Statistical significance was calculated utilizing three biological replicates for each exposure condition and genotype. Statistical significance was determined using a student’s unpaired t-test.

### 2.7 RNA Next-Generation Sequencing

Total RNA was isolated from liver tissue. Samples were placed in 1 mL of Trizol reagent (Invitrogen, CA), then homogenized with a Tissue-Tearor homogenizer (BioSpec, Bartlesville, OK). RNA isolation was performed using manufacturer’s protocol (catalog no. 15596026, Thermo Fisher Scientific). Isolated RNA samples were treated with DNase I, RNase-free (1 U/µL) (catalog no. EN0521, Thermo Scientific) following manufacturer’s instruction. Samples were then processed at The Center of Functional Genomics/University at Albany for differential expression with Poly-A selection and sequenced using an Illumina Nextseq500 (Illumina, Inc., San Diego, CA) at 75 million single-end reads per run. FASTQ files are available on NCBI Gene Expression Omnibus (GSE169155). Analysis was performed through bash and differential expression was analyzed using R Studio v1.3.1093 (Team 2020)(Boston, MA). Enhanced volcano plots were created using DESeq2 package (Love et al. 2014) and EnhancedVolcano (Blighe, Rana, and Lewis 2020) with a fold change (FC) cutoff of 1.0 and a pCutoff of 0.05. Gene count plots of epitranscriptomic writers were created using DeSeq2 and ggplot2 (Wickham 2016) packages. Metascape analysis (Zhou et al. 2019) was performed with upregulated genes of a log_2_-fold change cut-off of 2.0 or greater and ensembl ID lists were uploaded respectively per comparison condition. Statistical significance was calculated utilizing three biological replicates for each exposure condition and genotype, and was determined using student’s unpaired t-test.

## Results

### 3.1 *Alkbh8^Def^* mice show hypersensitivity to APAP toxicity and dysregulated selenoprotein expression in the liver

APAP can be bioactivated by a cytochrome P450 enzyme, Cyp2e1, to catalyze the formation of the reactive metabolite NAPQI (**Fig. 1A**). The response to toxic levels of APAP (600 mg/kg) was studied in 8–12-week-old C57BL/6 male WT and *Alkbh8^Def^* mice. Stress biomarkers indicative of liver damage and oxidative stress were measured in serum derived from whole blood and liver tissue, respectively. Alanine aminotransferase (ALT) levels were significantly higher in *Alkbh8^Def^* livers 6 hr after injection of APAP, relative to APAP exposed WT livers (**Fig. 1B**). The lipid peroxidation product 8-isoprostane is an *in vivo* biomarker of oxidative stress (Montuschi, Barnes, and Roberts 2004). For both saline and 6 hr APAP exposure conditions for *Alkbh8^Def^* liver tissue there was a slight increase in 8-isoprostane levels compared to WT samples (**Fig.1C**), but it did not reach a significant level (p = 0.33). Glutathione peroxidase 3 (Gpx3) is an extracellular antioxidant enzyme that aids in scavenging hydrogen peroxide as well as other hydroperoxides (Takebe et al. 2002). In response to APAP, there is a significant increase (p = 0.02) in Gpx3 expression in WT compared to its saline control, and this increased response was not observed in *Alkbh8^Def^* liver tissue (**Fig.1D**).

**Figure 1.**
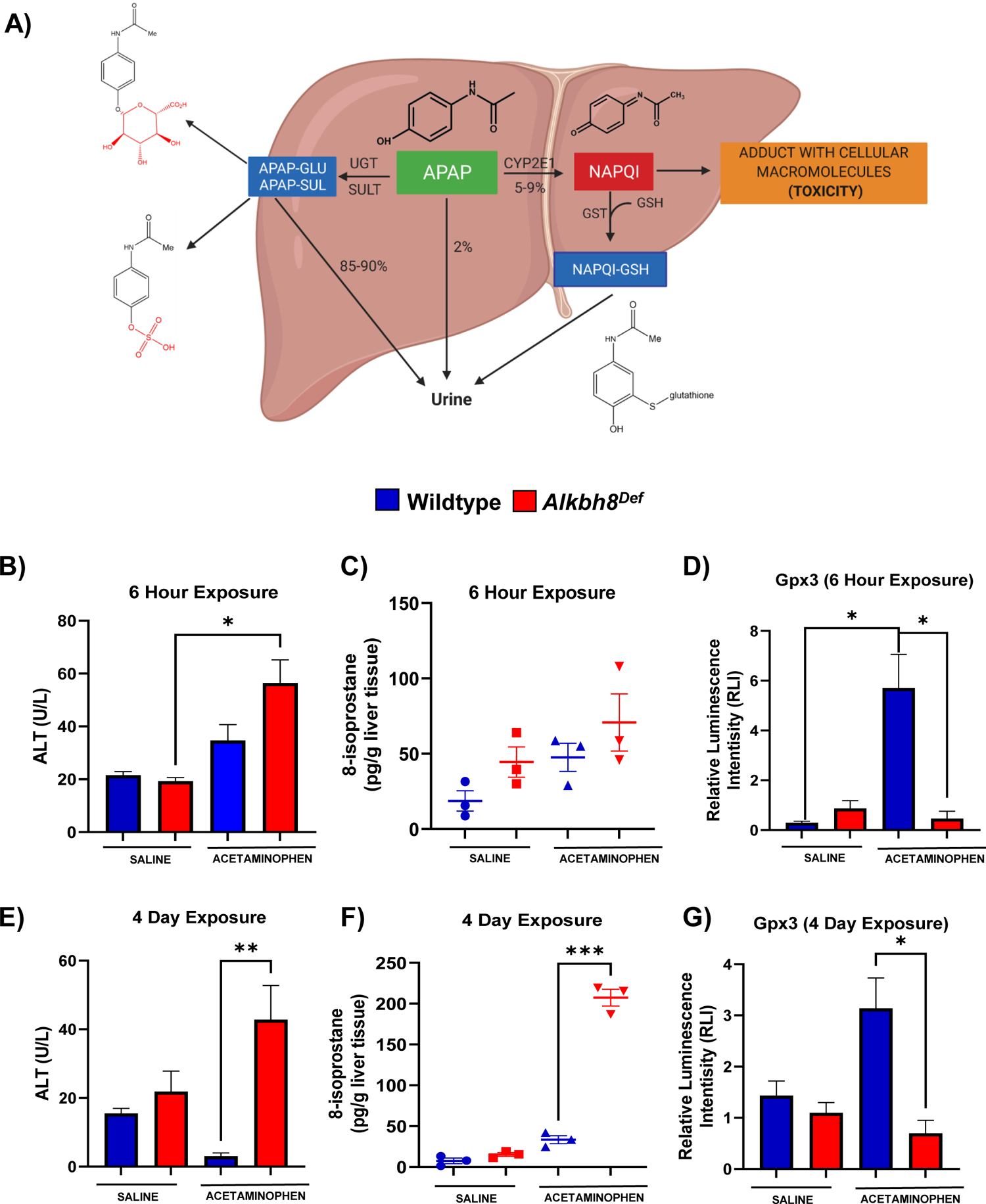
Writer deficient mouse livers have increased sensitivity to APAP induced stress and display decreased selenoprotein levels. (A) APAP can be converted into a non-toxic form by UDP-glucuronosyltransferases (UGT) or a toxic form by cytochrome P450 enzyme (CYP2E1) to generate the reactive intermediate metabolite N-actyl-p-benzoquinone imine (NAPQI). Male C57Bl/6 mice WT (blue) and *Alkbh8^Def^* (red) mice (8-12 weeks old) were exposed to a single 600 mg/kg dose of APAP (B-D) or daily 600 mg/kg doses of APAP over 4 days (E-G), and tissue/blood was harvested. (B, E) ALT and (C, F) 8-isoprostane levels were determined in blood and liver samples, respectively. (D, G) Gpx3 protein levels in the liver were evaluated using the ProteinSimple WES system. Statistical significance of biological replicates (N = 3) was measured by an unpaired t-test with (*p < 0.05, **p < 0.01, *** p < 0.001).

Daily injections over 4 days were used to study chronic APAP use in the WT and *Alkbh8^Def^* mice. ALT (p = 0.004) and 8-isprostane levels (p = 0.0001) were measured and found to be significantly increased in the *Alkbh8^Def^* mice after APAP injection compared to WT APAP (**Fig. 1E-F**). Gpx3 expression levels in WT and *Alkbh8^Def^* livers were also significantly (p = 0.02) different after 4 days of APAP exposure (**Fig. 1G**). Similar to 6-hour data, there was an APAP-induced increase in Gpx3 in WT livers, which was not observed in the *Alkbh8^Def^* mice. Our ALT, 8-isoprostane and Gpx3 data support the idea that writer deficient livers are experiencing significant APAP induced stress and defective selenoprotein synthesis.

### 3.2 Next-generation sequencing analysis reveals similar transcriptional responses to a single dose of APAP between WT and *Alkbh8^Def^* livers

We next analyzed gene expression using mRNA-seq of livers from saline and APAP-injected mice. Differential transcript expression was first visualized using Enhanced Volcano plots, with a p_adj._ values ≤ 0.05, from studies using a single injection of APAP (6 hours). WT APAP vs. WT saline (**Fig. 2A**) and *Alkbh8^De**f**^* APAP vs. *Alkbh8^Def^* saline (**Fig. 2C**) both show an extensive change in gene expression, with many transcripts significantly increase in response to APAP. Enrichment of biological pathways in significantly up-regulated genes (log_2_-fold change > 2.0 and padj. values ≤ 0.05) demonstrated that the following similar pathways were up-regulated in both WT (**Fig. 2B**) and *Alkbh8^Def^* (**Fig. 2D**) livers; MAPK signaling pathway, positive regulation of cell death, fat cell differentiation, osteoclast differentiation, response to growth factor, and regulation of neuron death. In general, both genotypes regulate groups of transcripts linked to stress responses. A comparison of mRNA seq results (**Fig. 3A**) further supports the idea that the transcriptional response to APAP is very similar in WT and *Alkbh8^Def^* livers where only 202 out of 55,487 measured transcripts were significantly different (p < 0.05) between the two genotypes. Transcriptional upregulated genes that had the largest significant difference in fold change include: odorant binding protein 2A (*Obp2a)* which aids in binding and transporting molecules with high affinity for aldehydes and large fatty acids (Briand et al. 2002), transmembrane protein 267 (*Tmeme267)* which present reported function involves protein binding (Strausberg et al. 2002) and a member of the MYC Proto-Oncogene (*myc)* family that is highly expressed in the brain (*Bmyc)* and functions as a transcriptional regulator. We also report similar gene transcript profiles when comparing WT and *Alkbh8^Def^* under saline control conditions (**Fig. S1**). The similarities in the transcriptional responses to APAP between the two genotypes was expected, as Alkbh8, along with its accessory protein Trm112, post transcriptionally regulate gene expression by modifying tRNA^Sec^ (**Fig. 3B-D**) and other tRNAs. Alkbh8 can modify tRNA^Sec^, which is used in stop codon recoding, to drive the translation of 25 selenocysteine containing proteins in the *Mus musculus* genome (**Fig. 3D**). Other regulatory components tied to stop codon recoding include the cis Sec insertion sequence (SECIS), a stem-loop structure located in coding regions and in 3’UTRs of eukaryotic genes (Low & Berry, 1996; Berry, J et al., 1991).

**Figure 2.**
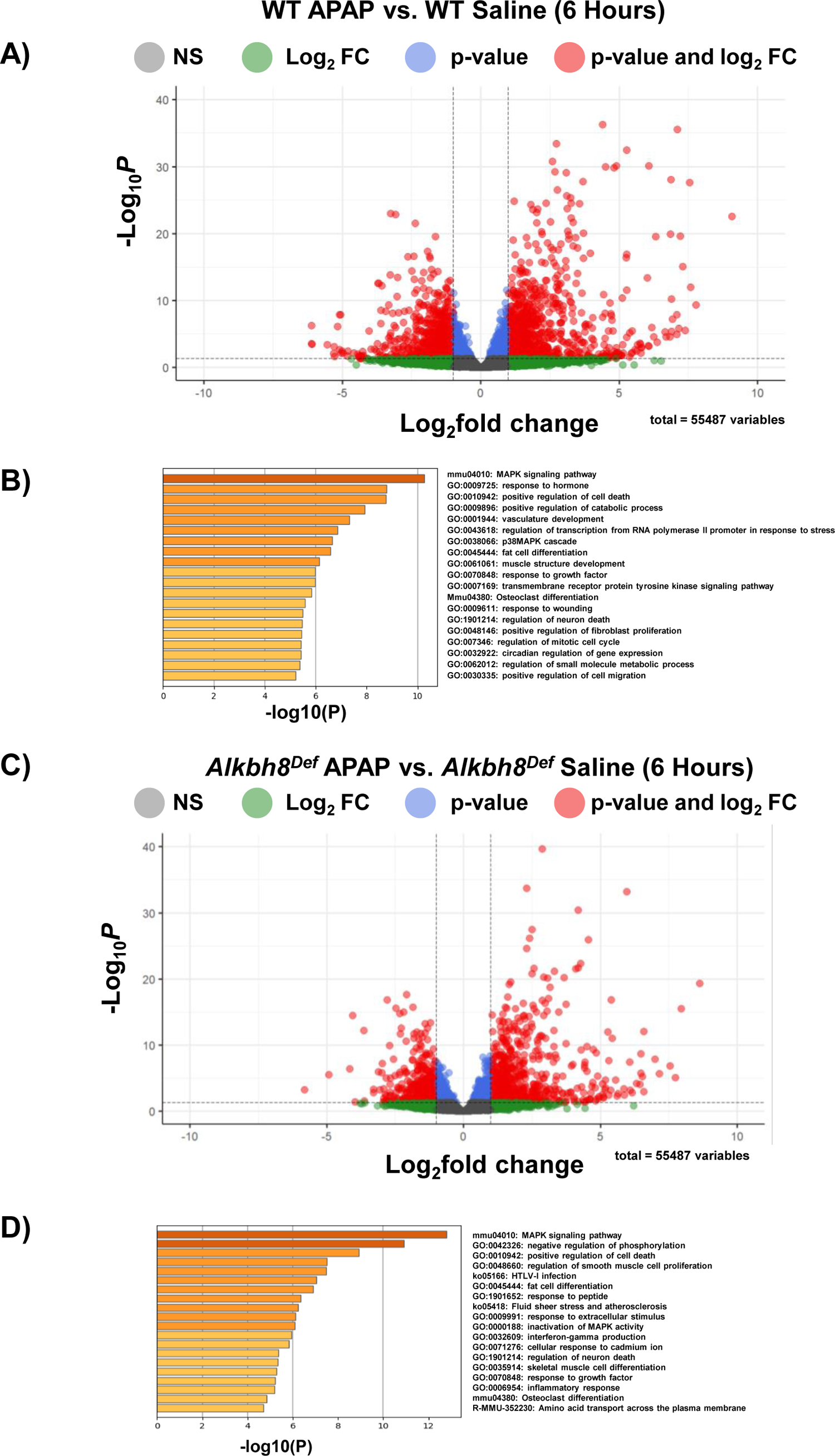
Next-generating sequencing analysis of the transcriptional response to APAP. WT and *Alkbh8^Def^* mice (N = 3) were left untreated or exposed to a single dose of 600 mg/kg APAP and livers were harvested after 6 hours and RNA was purified and subject to mRNA-seq. Enhanced volcano plots for each of the comparisons and Metascape analysis were performed for log_2_FC > 2.0 for **(A-B)** WT APAP vs. WT Saline and **(C-D)** *Alkbh8^Def^* APAP vs. *Alkbh8^Def^* Saline.

**Figure 3.**
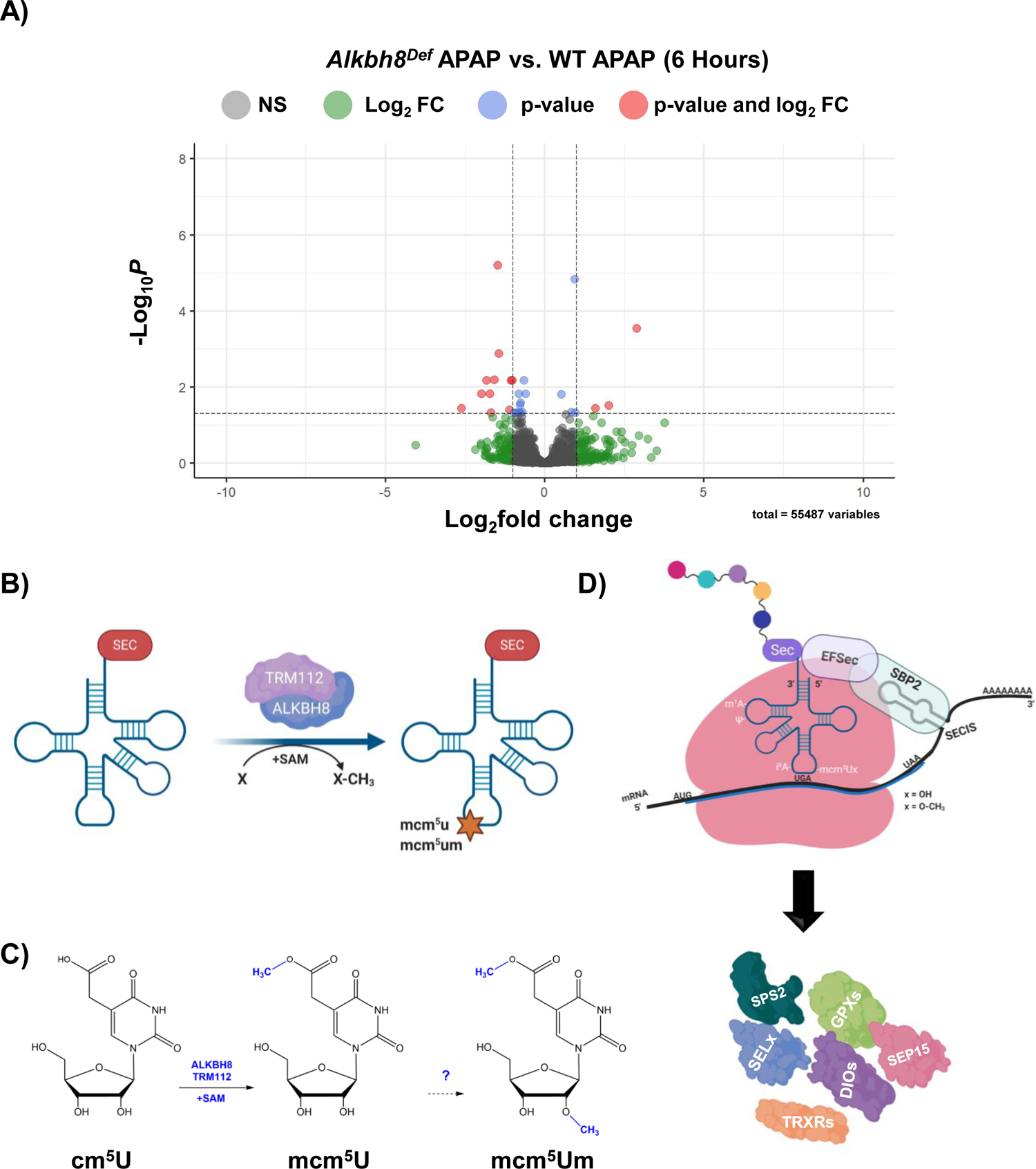
WT and *Alkbh8^Def^* livers have a similar transcriptional response to a single dose of APAP. mRNA-seq data for **(A)** *Alkbh8^Def^* APAP vs. WT APAP was compared using an enhanced volcano plot. **(B)** Post-transcriptional regulation of translation by ALKBH8 and epitranscriptomic marks. Alkbh8 requires the small accessory protein Trm112 and S-adenosylmethionine (SAM) to modify the wobble position of tRNA^Sec^ **(C)** from cm^5^U to mcm^5^U, which is then further modified to the ribose-methylated derivative mcm^5^Um. **(D)** Stop-codon recoding utilizes mcm^5^U and mcm^5^Um in tRNA^Sec^ to decode UGA to produce selenoproteins. In addition to Alkbh8-catlyzed epitranscriptomic marks, elongation factors, a selenocysteine insertion sequence (SECIS), accessory proteins and an internal UGA stop codon are used to promote the translation of selenocysteine containing proteins.

### 3.3 The epitranscriptome is extensively regulated in response to APAP and disrupted in ***Alkbh8^Def^ livers*.**

We next analyzed for differences in post transcriptional regulation by quantifying epitranscriptomic marks in RNA from WT and *Alkbh8^De**f**^* livers. tRNA^Sec^ possesses four different modifications (**Fig. 4A**) that are catalyzed by either Alkbh8 (mcm^5^U), Trmt61A/Trm6 (m^1^A), Pus4 (Ψ), and Trit1 (i^6^A). We used LC-MS/MS based approaches to show that all four RNA modifications are increased in WT livers in response to APAP, with these increases absent in *Alkbh8^De**f**^* livers. The WT livers had significantly (p = 0.04) increased the concentration of the m^1^A modification from 55.51 pg/µL to 85.54 pg/µL, whereas the *Alkbh8^Def^* liver showed a decreased concentration from saline of 49.94 pg/µL to 38.48 pg/µL after APAP exposure. Similarly, Ψ was significantly (p = 0.002) increased in WT saline (49.12 pg/µL) to WT APAP (60.76 pg/µL) conditions and showed a decreased concentration in *Alkbh8^De**f**^* saline (51.10 pg/µL) when compared to *Alkbh8^De**f**^* APAP (35.71 pg/µL) exposed liver tissue. The i^6^A modification was also dysregulated in the *Alkbh8^De**f**^* mice, where we measured a decreased concentration of 0.54 pg/µL in saline verse 0.42 pg/µL for APAP exposure. WT liver tissue was measured to significantly (p = 0.04) increase i^6^A modification from 0.64 pg/µL in saline conditions to 1.25 pg/µL after APAP exposure. Finally, the Alkbh8 dependent mcm^5^U modification was measured to have minimal change in concentration in the *Alkbh8^De**f**^* liver tissue between saline (0.0036 pg/µL) and APAP (0.0035 pg/µL) conditions, unlike the WT liver tissue where we measured a significant (p = 0.01) increase from 0.004 pg/µL to 0.045 pg/µL in saline and APAP conditions, respectively. The failure to increase epitranscriptomic marks in tRNA^Sec^ in response to APAP in *Alkbh8^De**f**^* liver tissue predicts that selenoproteins levels would be affected. We quantified Gpx1 and Trxr2 levels (**Fig. 4C**) and similar to Gpx3 (**Fig. 1D**), Gpx1 and Trxr2 were increased in WT livers exposed to APAP with Trxr2 being significant (p = 0.002); this stress induced increase in protein levels was absent in *Alkbh8^De**f**^* livers. Gpx4, Trxr1 and selenoprotein S (SelS) also failed to respond to APAP treatment in *Alkbh8^De**f**^* animals **(Supplemental Fig. S4**).

**Figure 4.**
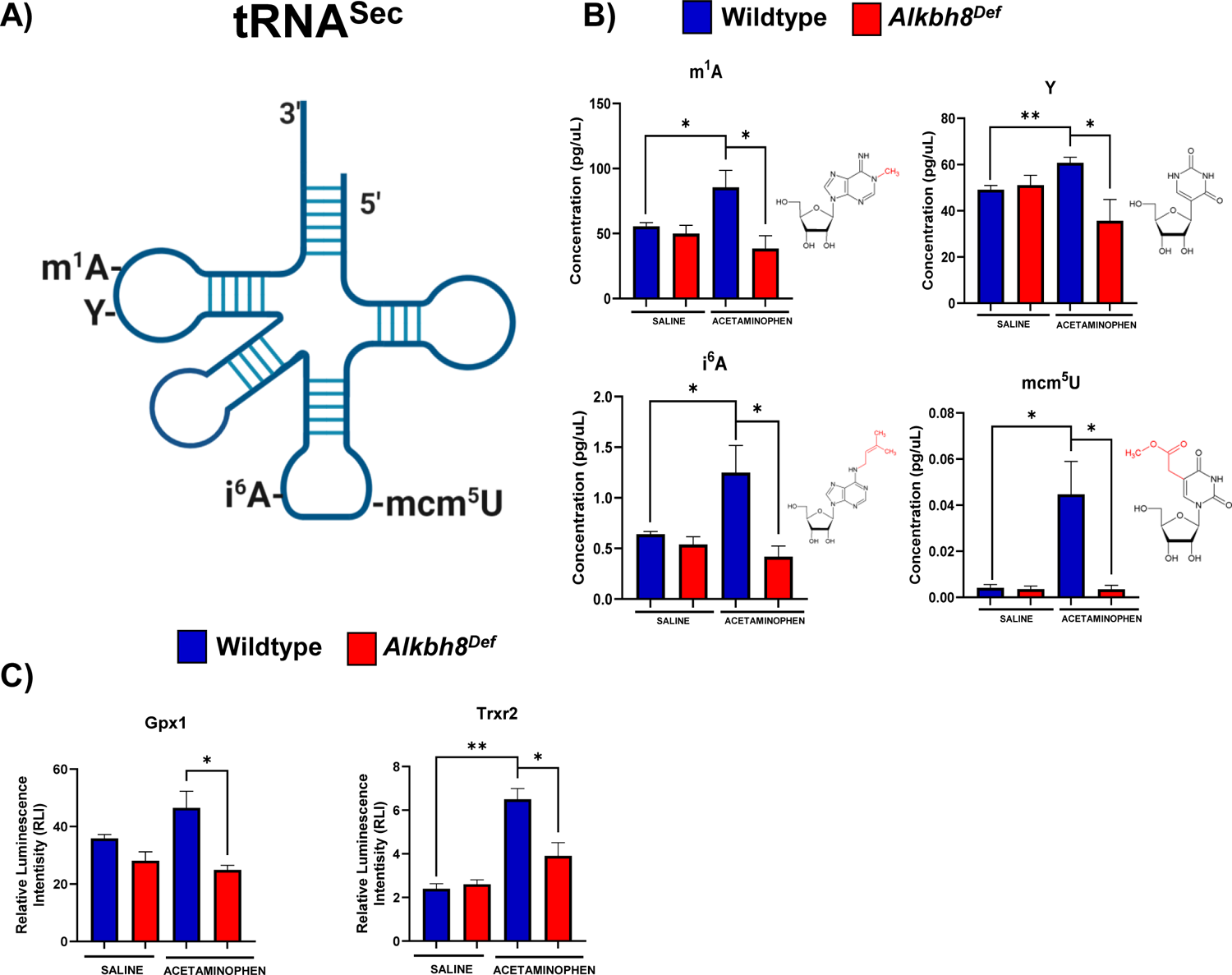
Multiple RNA modifications on tRNA for selenocysteine are differentially regulated in WT and *Alkbh8^Def^* livers in response to a single dose of APAP. (A) Epitranscriptomic marks found on tRNA^Sec^. **(B - C)** WT and *Alkbh8^Def^* mice (N=3) were exposed to a single 600 mg/kg dose of APAP and livers were harvested 6 hours after dosing. **(B)** Modifications that occur throughout tRNA^Sec^ were measured using LC-MS/MS. **(C)** Selenoprotein levels were evaluated using the ProteinSimple WES system. Statistical significance of biological replicates (N = 3) was determined using an unpaired t-test with (*p < 0.05, **p < 0.01, *** p < 0.001).

We measured 37 tRNA modifications using LC-MS/MS and compared their levels, relative to WT saline, using a heat map (**Fig. 5A-B**). The WT response to APAP resulted in 31 total increased modifications with 22 of the group being significant (**Table 1**) whereas the *Alkbh8^Def^* livers have 32 modifications decreased with 13 significant modifications. All 17 modifications that are decreased in *Alkbh8^Def^* saline control conditions, relative to WT saline are significant (**Table 1**). Individual bar graphs of tRNA modifications from the heat map show mean values with standard error around the mean in WT and *Alkbh8^Def^* liver tissue under APAP exposure conditions (**Fig. 5B**). Other modifications are shown in (**Supplemental Fig. S5**). The tRNA modifications that were the most downregulated in the *Alkbh8^Def^* APAP samples were; T (−6.5 log_2_fold change, p < 0.05), mnm^5^U (−6.4 log_2_fold change, p < 0.05) and mcm^5^U (−3.7 log_2_fold change, p < 0.05). Our epitranscriptomic data provides strong support for the idea that a single acute toxic dose of APAP promotes significant changes in many tRNA modifications. Below we explore whether chronic APAP exposure promote changes to the epitranscriptome.

**Figure 5.**
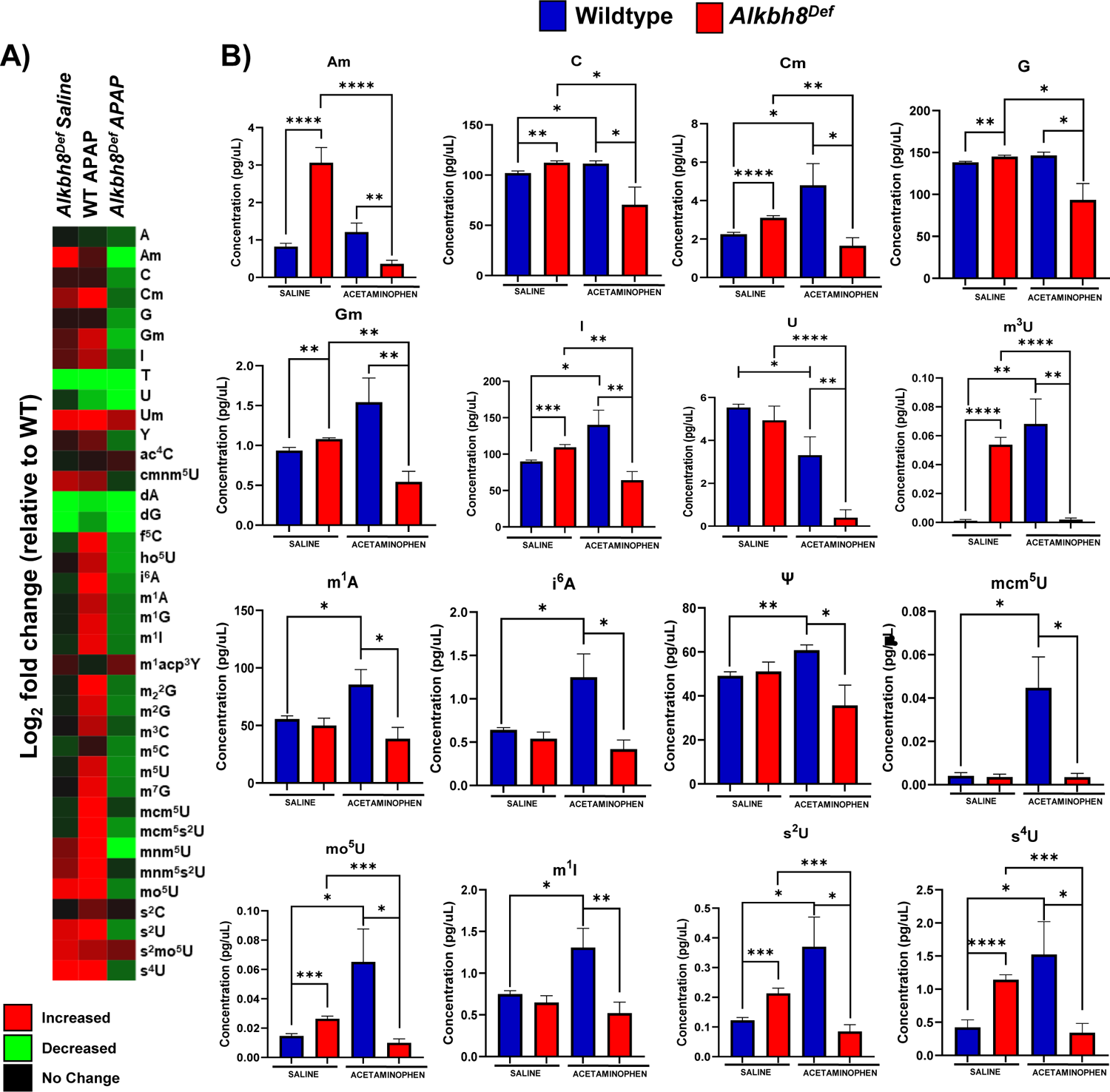
The epitranscriptome is differentially regulated in WT and *Alkbh8^Def^* livers in response to a single dose of APAP. (A) 37 epitranscriptome marks were measured with LC-MS/MS and the log_2_-fold change was normalized relative to WT saline data and shown as a heat map. **(B)** Data for each modification under the four conditions was compared for statistical significance, with the average of biological replicates (N = 3) using an unpaired t-test with (*p < 0.05, **p < 0.01, *** p < 0.001).

**Table 1.** Measured epitranscriptomic marks in mouse liver tissue after APAP (6 hours). Calculations for each epitranscriptomic mark and comparisons between WT and *Alkbh8^Def^* liver tissue post 6-hour exposure to 600 mg/kg of APAP with reported statistical significance of biological replicates (N = 3) measured by an unpaired t-test.

| tRNA Modification | *Alkbh8*<sup>Def</sup> Saline vs. WT Saline, Log<sub>2</sub> Fold Change | WT APAP vs. WT Saline, Log<sub>2</sub> Fold Change | *Alkbh8*<sup>Def</sup> APAP vs. WT Saline, Log<sub>2</sub> Fold Change | *Alkbh8*<sup>Def</sup> APAP vs. WT APAP, Log<sub>2</sub> Fold Change | *Alkbh8*<sup>Def</sup> APAP vs. *Alkbh8*<sup>Def</sup> Saline, Log<sub>2</sub> Fold Change | WT APAP vs. WT Saline p-Value | *Alkbh8*<sup>Def</sup> Saline vs WT Saline p-Value | *Alkbh8*<sup>Def</sup> APAP vs. WT APAP p-Value | *Alkbh8*<sup>Def</sup> APAP vs. *Alkbh8*<sup>Def</sup> Saline p-Value |
| --- | --- | --- | --- | --- | --- | --- | --- | --- | --- |
| A | 0.0 | -0.1 | -0.3 | -0.2 | -0.28 | 0.0340 | 0.0470 | 0.0460 | 0.0037 |
| Am | 1.6 | 0.2 | -1.5 | -1.7 | -3.08 | 0.0700 | 0.0001 | 0.0022 | 0.000004 |
| C | 0.2 | 0.1 | -0.5 | -0.7 | -0.67 | 0.0920 | 0.0005 | 0.0180 | 0.0160 |
| Cm | 0.6 | 1.2 | -0.4 | -1.5 | -0.91 | 0.3010 | 0.0001 | 0.0100 | 0.0020 |
| G | 0.1 | 0.1 | -0.6 | -0.6 | -0.63 | 0.0680 | 0.0011 | 0.0084 | 0.0088 |
| Gm | 0.3 | 0.8 | -0.7 | -1.5 | -1.00 | 0.3210 | 0.0032 | 0.0042 | 0.0006 |
| I | 0.3 | 0.7 | -0.5 | -1.1 | -0.77 | 0.2840 | 0.0001 | 0.0024 | 0.0011 |
| T | -6.7 | -1.2 | -7.7 | -6.5 | -1.02 | 0.0000 | 0.0003 | 0.0440 | 0.1250 |
| U | -0.1 | -0.7 | -3.8 | -3.1 | -3.65 | 0.3060 | 0.2180 | 0.0034 | 0.00001 |
| Um | 0.9 | 1.4 | 0.6 | -0.8 | -0.30 | 0.3440 | 0.1110 | 0.3100 | 0.3350 |
| Y | 0.1 | 0.4 | -0.4 | -0.8 | -0.52 | 0.0046 | 0.2080 | 0.0090 | 0.0740 |
| ac<sup>4</sup> C | -0.1 | 0.1 | 0.2 | 0.1 | 0.22 | 0.0750 | 0.4190 | 0.0036 | 0.1100 |
| cmnm<sup>5</sup> U | 0.7 | 0.5 | -0.2 | -0.7 | -0.86 | 0.1700 | 0.0240 | 0.0620 | 0.0110 |
| dA | -4.6 | -0.9 | -5.5 | -4.6 | -0.90 | 2.10E-13 | 0.0000004 | 0.0210 | 0.1240 |
| dG | -4.8 | -0.6 | -5.5 | -5.0 | -0.76 | 0.0000001 | 0.00002 | 0.0240 | 0.1340 |
| f<sup>5</sup> C | -0.2 | 1.0 | -0.6 | -1.6 | -0.40 | 0.0190 | 0.3690 | 0.0190 | 0.3130 |
| ho<sup>5</sup> U | 0.0 | 0.7 | -0.6 | -1.4 | -0.67 | 0.0110 | 0.4720 | 0.0230 | 0.1750 |
| i<sup>6</sup> A | -0.2 | 1.1 | -0.5 | -1.6 | -0.36 | 0.0001 | 0.2530 | 0.0055 | 0.1870 |
| m<sup>1</sup> A | -0.1 | 0.7 | -0.4 | -1.2 | -0.38 | 0.0001 | 0.4010 | 0.0053 | 0.1710 |
| m<sup>1</sup> G | -0.1 | 0.9 | -0.4 | -1.3 | -0.38 | 0.0001 | 0.4000 | 0.0060 | 0.1730 |
| m<sup>1</sup> I | -0.1 | 0.9 | -0.4 | -1.3 | -0.31 | 0.0001 | 0.3120 | 0.0045 | 0.2140 |
| m<sup>1</sup> acp<sup>3</sup> ψU | 0.2 | -0.1 | 0.4 | 0.4 | 0.17 | 0.0350 | 0.2690 | 0.0490 | 0.1690 |
| m<sup>22</sup> G | -0.1 | 1.2 | -0.4 | -1.6 | -0.34 | 0.0036 | 0.4110 | 0.0095 | 0.1890 |
| m<sup>2</sup> G | -0.1 | 0.8 | -0.5 | -1.3 | -0.39 | 0.0001 | 0.3830 | 0.0058 | 0.1660 |
| m<sup>3</sup> C | 0.0 | 0.7 | -0.3 | -0.9 | -0.28 | 0.0001 | 0.4760 | 0.0100 | 0.2390 |
| m<sup>5</sup> C | -0.2 | 0.1 | -0.5 | -0.6 | -0.26 | 0.0002 | 0.1770 | 0.0340 | 0.2620 |
| $m^5 U$ | -0.1 | 0.8 | -0.5 | -1.3 | -0.44 | 0.0001 | 0.3940 | 0.0042 | 0.1340 |
| $m^7 G$ | 0.0 | 0.9 | -0.5 | -1.4 | -0.46 | 0.0003 | 0.4990 | 0.0046 | 0.1220 |
| $mcm^5 U$ | -0.1 | 3.5 | -0.2 | -3.7 | -0.04 | 0.0370 | 0.4310 | 0.0056 | 0.4820 |
| $mcm^5 s^2 U$ | -0.1 | 1.0 | -0.5 | -1.6 | -0.41 | 0.0001 | 0.2990 | 0.0066 | 0.1500 |
| $mnm^5 U$ | 0.4 | 4.7 | -1.7 | -6.4 | -2.14 | 0.0390 | 0.0150 | 0.0290 | 0.000001 |
| $mnm^5 s^2 U$ | 0.5 | 3.1 | -0.1 | -3.2 | -0.62 | 0.3430 | 0.0440 | 0.0290 | 0.0500 |
| $mo^5 U$ | 0.9 | 2.3 | -0.5 | -2.7 | -1.41 | 0.2330 | 0.00004 | 0.0130 | 0.0001 |
| $s^2 C$ | 0.0 | 0.4 | 0.0 | -0.3 | 0.04 | 0.1170 | 0.4970 | 0.1220 | 0.4510 |
| $s^2 U$ | 0.8 | 1.6 | -0.5 | -2.1 | -1.33 | 0.4040 | 0.0001 | 0.0067 | 0.0002 |
| $s^2 mo^5 U$ | 0.9 | 0.6 | 0.4 | -0.2 | -0.44 | 0.3310 | 0.0260 | 0.2360 | 0.1030 |
| $s^4 U$ | 1.4 | 1.8 | -0.3 | -2.2 | -1.74 | 0.0007 | 0.00002 | 0.0180 | 0.0001 |

### 3.4 The *Alkbh8* writer deficient mice are sensitive to daily APAP doses and respond with altered gene expression

We analyzed RNA from mice on the 4^th^ day of daily APAP treatments. mRNA sequencing experiments revealed many (N=3) differences in gene regulation between WT and *Alkbh8^Def^* in the response to APAP toxicity at the 4^th^ day. The comparison of mRNA-seq data from WT APAP vs. WT saline identified significantly upregulated genes (p ≤ 0.05) in pathways related to response to stillbenoid, inflammatory response and unsaturated fatty acid metabolic process (**Fig. 6B**). The *Alkbh8^Def^* APAP vs. *Alkbh8^Def^* Saline mRNA-seq comparison shows significantly (p ≤ 0.05) upregulated genes that are enriched for pathways related to; mitotic cell cycle process, cellular response to DNA damage stimulus, and positive regulation of cell cycle (**Fig. 6C**). A comparison of saline controls between WT and *Alkbh8^Def^* show very few differences in transcript levels at the 4-day time point **(Supplemental Fig. S2**). However, unlike the single APAP dose, the multiple daily doses of APAP promoted many differences in gene regulation between WT and the *Alkbh8^Def^* liver tissue **(Supplemental Fig. 3A**). Pathways specific to cell division, regulation of cell cycle process and cellular response to DNA damage stimulus were significantly (p ≤ 0.05) enriched in *Alkbh8* APAP vs. WT APAP comparison **(Supplemental Fig. 3B**), supporting the idea that there is more stress and DNA damage in *Alkbh8^Def^* livers compared to WT. The same selenoproteins that were measured for the 6-hour timepoint were quantitated after 4 days of daily doses of APAP in liver tissue. Gpx1 expression in the WT mice show an increase in the APAP exposure group, while the *Alkbh8^Def^* liver tissue shows decreased Gpx1. Gpx4 was elevated in the WT as well after APAP, while there was a significant decrease in the expression of Gpx4 in the *Alkbh8^Def^* liver. Both TrxR1 and TrxR2 protein levels were increased in WT APAP samples in response to saline, which was not observed in *Alkbh8^Def^* livers (**Fig. 6D**).

**Figure 6.**
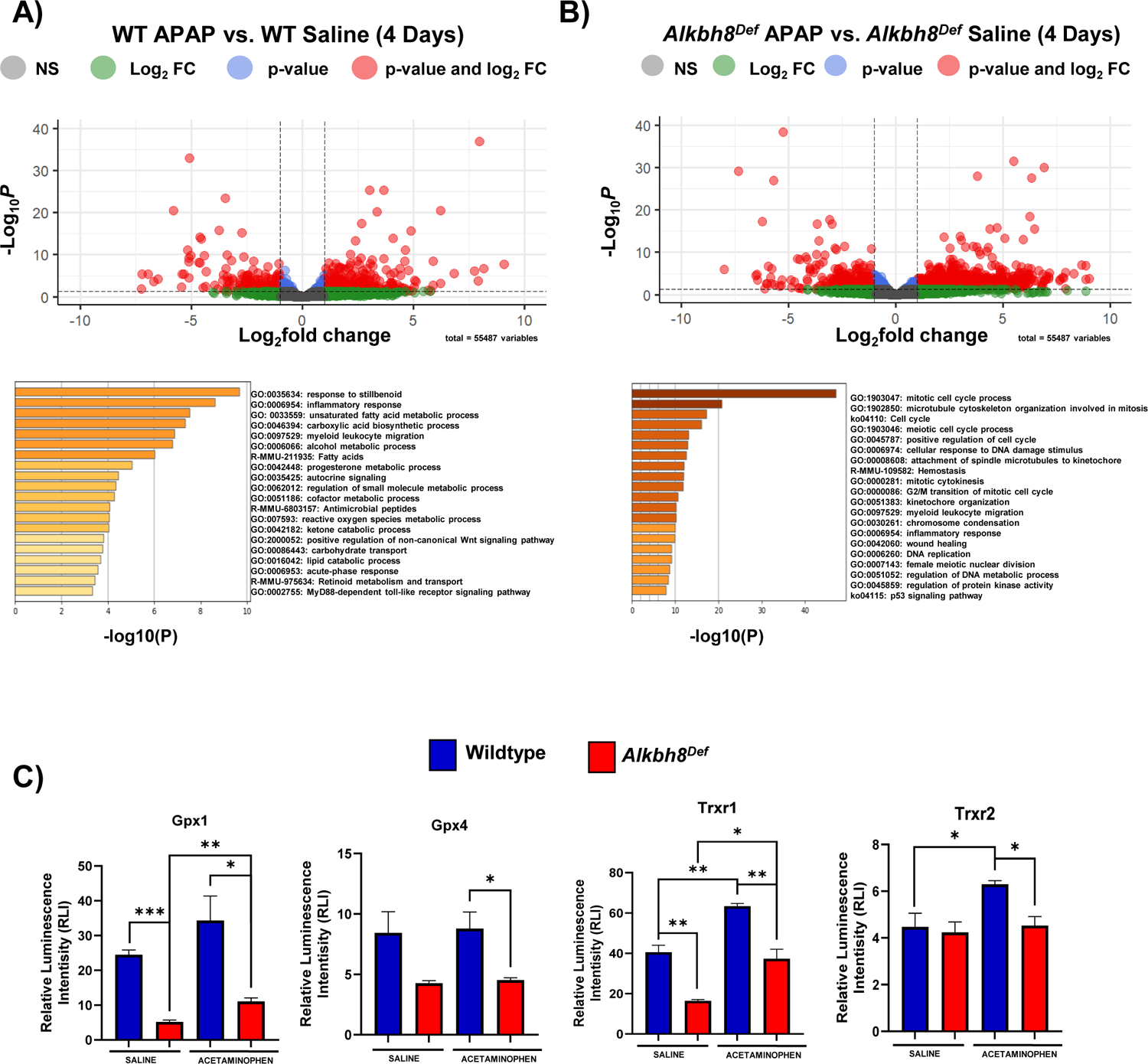
Writer deficient mice are sensitive to daily APAP, with altered gene expression and response due to altered epitranscriptomic response? WT (N = 3) and *Alkbh8^Def^* (N = 3) mice were IP injected daily with 600 mg/kg of APAP. (A-C) Tissue was harvested at day 4 and RNA and protein were purified for analysis. (A-B) Enhanced volcano plots were generated for each of the comparisons and Metascape analysis was performed for log_2_FC > 2.0. (A) WT APAP vs. WT Saline and (B) *Alkbh8^Def^* APAP vs. *Alkbh8^Def^* Saline. (C) Selenoprotein levels were evaluated using the ProteinSimple WES system. Statistical significance of biological replicates (N = 3) was determined using an unpaired t-test with (*p < 0.05, **p < 0.01, *** p < 0.001).

Results of LC-MS/MS analysis confirmed the disrupted response of most tRNA^Sec^ and wobble U related modifications in a chronic timeline. In response to APAP the WT liver tissue showed an increase in 6 significant (p ≤ 0.05) modifications and 2 significant (p ≤ 0.05) decreased modifications relative to WT saline. The *Alkbh8^Def^* mice had 9 significant (p ≤ 0.05) modifications decreased after APAP exposure and increased 5 significant (p ≤ 0.05) modifications relative to *Alkbh8^Def^* saline. Under APAP exposure conditions 3 significant (p ≤ 0.05) increased modifications and 12 significant (p ≤ 0.05) decreased modifications were revealed in *Alkbh8^Def^* liver tissue relative to WT. tRNA^Sec^ related modifications revealed a dynamic change in the epitranscriptomic response after four daily doses of APAP. The modification that had a similar response to acute conditions was m^1^A. The WT liver tissue measured a significant (p = 0.01) increase after APAP exposure unlike the *Alkbh8^Def^* liver tissue which was measured to have a significant (p = 0.00003) decrease when comparing saline to APAP conditions. Many wobble uridine modifications were measured to be dysregulated in the *Alkbh8^Def^* liver tissue compared to WT after daily APAP exposure including: m^5^U, mcm^5^s^2^U, mnm^5^U, mo^5^U, mnm^5^s^2^U, and s^4^U. It is interesting to note that both s^2^U and s^2^mo^5^U concentration were measured to be increased after daily APAP exposure in *Alkbh8^Def^* mice compared to WT. Adenosine-based modifications m^1^A, and m^6^A were also measured to be dysregulated in the *Alkbh8^Def^* liver tissue compared to WT after daily APAP exposure. All measured modifications within the chronic timeline are shown in **(Supplemental Fig. 7**) and (**Supplemental Table 1**).

**Figure 7.**
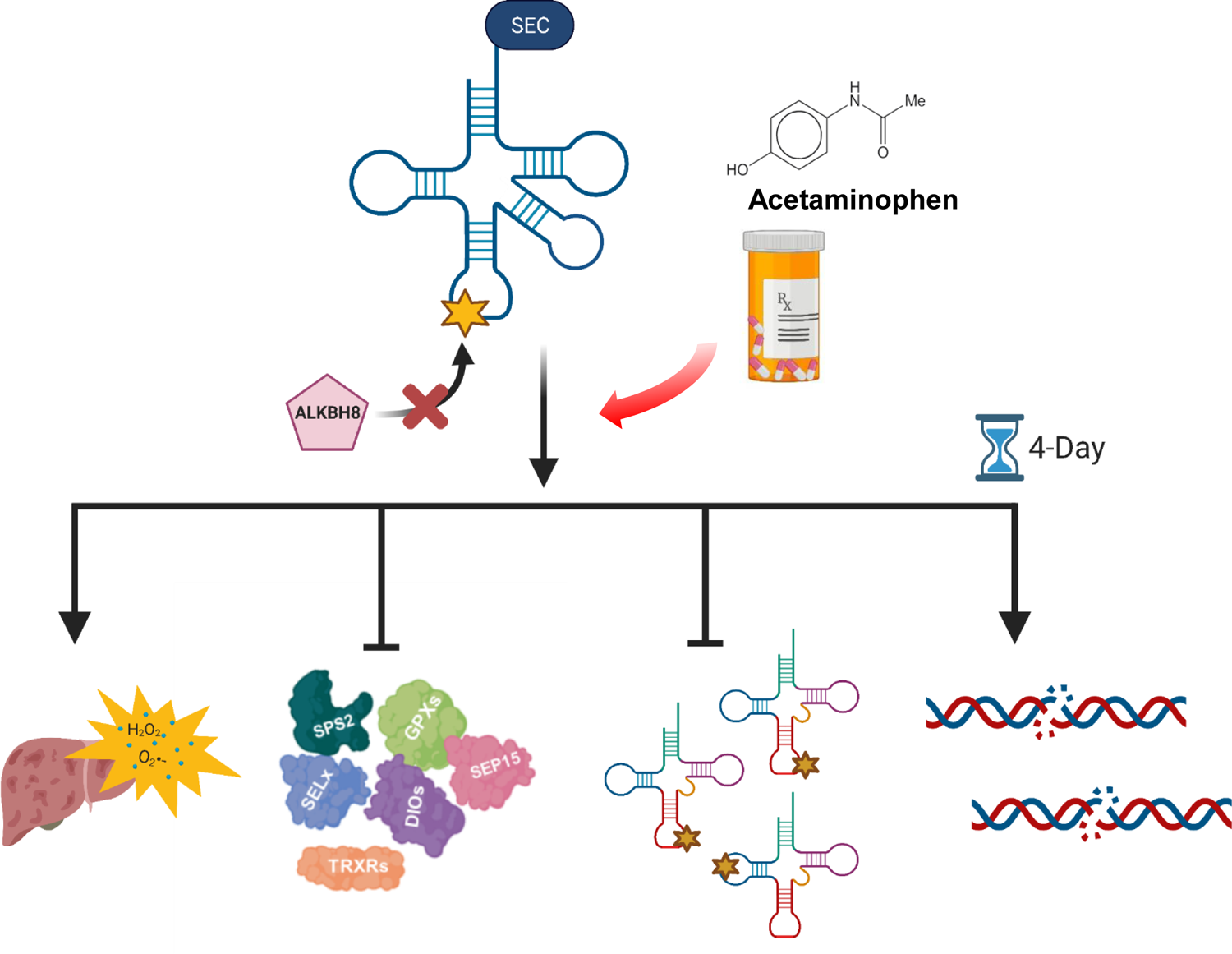
Model of APAP sensitivity in writer and modification dysregulated Alkbh8^Def^ livers. The deficiency of A*lkbh8* uncovers the protective role against oxidative stress-induced APAP toxicity. *Alkbh8^Def^* mice experienced higher instances of oxidative stress, liver damage and overall decreased survival. The deficiency of *Alkbh8* results in the dysregulation of the epitranscriptome and selenoprotein expression. Chronic APAP exposure results in an increased transcriptional expression of DNA repair and DNA damage response.

## Discussion

Hepatoxic effects of APAP toxicity have been widely reported which include oxidative stress, impaired mitochondrial function, lipid peroxidation, nuclear DNA fragmentation and necrotic cell death (Du, Ramachandran, and Jaeschke 2016; Jaeschke and Ramachandran 2018; Mcgill et al. 2012; Mossanen and Tacke 2015; Yan et al. 2010). ROS production from the bioactivation of APAP can promote DNA and protein damage (Borude, Bhushan, and Apte 2018; McGill et al. 2014; Ramachandran and Jaeschke 2017). The *Alkbh8^Def^* mice were unable to respond to the same stress load as their WT counterparts, and our work supports the idea that the epitranscriptomic writer, Alkbh8, has a protective role against APAP toxicity. The protection comes at the levels of tRNA modification and translation, with Alkbh8 catalyzing tRNA modifications and translationally regulating the production of stress response proteins. Wobble U writers in yeast, mice and humans have previously been shown to play key roles in responding to stress by enhancing the translation of key ROS detoxification, DNA damage response and metabolic proteins (Chionh et al. 2016; Schaffrath and Leidel 2017; Villahermosa and Fleck 2017; Wilkinson, Cui, and He 2021). For example, deficiencies in the mcm^5^U writer tRNA methyltransferase 9 (Trm9) in yeast sensitize cells to DNA damaging agents and disrupt cell cycle transitions, with decreased translation of ribonucleotide reductase genes driving these phenotypes (Begley et al. 2007; Deng et al. 2015; Patil et al. 2012). Knockdown of human *ALKBH8* in HEK293 cells sensitize cells to ionizing radiation and disruption of mouse Alkbh8 sensitizes cells to the mitochondrial poison rotenone, due in part to the decrease in ROS detoxification enzymes (Endres et al. 2015b; Dragony Fu et al. 2010). Further it has been reported that there are increased markers of senescence in *Alkbh8^Def^* MEFs (Lee et al. 2020), highlighting a role for wobble U in regulating cell states linked to aging and disease. Notably, dysregulated expression of wobble U writers and their corresponding enzymes have also been linked to the etiology of cancer (Barbieri and Kouzarides 2020; Chellamuthu and Gray 2020; De Crécy-Lagard et al. 2019; Haruehanroengra et al. 2020; Popis, Blanco, and Frye 2016; C. Wang et al. 2021). Wobble U writers play important translational regulatory roles that protect normal cells and organs from stress, and our studies highlight that the wobble U writer Alkbh8 helps regulate the response to APAP toxicity.

### 4.1 Translational regulation is a key response to APAP

Protein kinase cascades are well described responses to APAP, as it activates the c-Jun-N-terminal kinase (JNK) through the sequential protein phosphorylation of mainly mitogen-activated protein kinases (MAPK)(Harrison et al. 2004; Johnson and Lapadat 2002). JNK is a critical member of the MAPK family and considered a control point between physiological and pathological status. The activation of JNK has shown to be important in biological processes including stress reaction, cellular differentiation, and apoptosis (Harrison et al. 2004; Johnson and Lapadat 2002). The activated JNK can translocate to the mitochondria and binds to an outer membrane protein known as Sab, which is also phosphorylated. JNK binding to Sab leads to further ROS generation and sustains the activation of JNK leading to the induction of the mitochondrial permeability transition (MPT) resulting in the mediation of hepatocyte necrosis (Win et al. 2011). *c-Myc* is induced by APAP and encompasses one of the most significant protein networks that are associated with liver injury (Beyer et al. 2007). Beyer *et al.,* (Beyer et al. 2007) presented gene ontology analysis of transcripts that significantly correlate with liver necrosis and the resulting biological pathways 6 hours post-APAP exposure included programmed cell death, response to wounding, and inflammatory response. These biological pathways are upregulated in both genotypes of mice after 6 hr APAP exposure when compared to their corresponding saline control. Our transcriptional studies support the idea that both WT and *Alkbh8^Def^* mice respond to APAP toxicity similarly at the mRNA level.

Our study is unique in that we describe a change in RNA modification regulated protein expression in response to APAP stress. Members of the Gpx and TrxR selenoprotein families are known to be critical for signaling and aiding in protection and reduction of harmful oxidants (Leonardi et al. 2019). Gpx and TrxR selenoproteins have also been shown to be key regulators of mitochondrial oxidative stress. Inhibition of these antioxidant enzymes with Auranofin in non-cancerous human Swan-71 cells resulted in reduction of cell viability, mitochondrial respiration, and increased oxidative stress. Selenium supplementation was able to restore Gpx and TrxR selenoprotein activity which subsequently decreased ROS production and increased cell viability (Radenkovic et al. 2017). These and other studies (Lopert, Day, and Patel 2012; Miyamoto et al. 2003) confirm the important role of Gpx and TrxR selenoproteins as key response regulators to mitochondrial oxidant stress. Furthermore, APAP hepatoxicity has been reported to result in the formation of the potent oxidant peroxynitrite from superoxide and nitric oxide (Hinson et al. 1998) which was subsequently observed to be predominately generated inside the mitochondria (Cover et al. 2005). Therefore, the mitochondria are a significant source of oxidant stress in response to APAP. Our study demonstrated under non-stressed conditions protein level expression of these selenoproteins are relatively similar and insignificantly different between WT and *Alkbh8^Def^*. However, after APAP exposure we see significant (p ≤ 0.05) differences, with WT increasing expression of selenoproteins and *Alkbh8^Def^* having no change or decreased expression in measured selenoproteins. Our identification of a translational response to APAP, specific to selenoproteins that most likely respond to mitochondrial stress, highlights the importance of protein synthesis-based responses to stress. Translational regulation is likely occurring in response to all pharmaceuticals that promote ROS stress, and studies in cancer models support the idea that this regulation will be extended to many types of protein families and include regulation of the 20 standard amino acids (Rapino et al. 2017, 2018).

### 4.2. RNA modifications are systematically and globally re-programmed in tRNA^Sec^ and other tRNAs in response to toxicants and now pharmaceuticals

tRNA modifications have been shown to be a critical part of epitranscriptomic reprogramming in response to stressors, chemicals and physiological states that have been shown in many organisms. (Chan et al. 2012; Endres, Dedon, and Begley 2015; Gu, Begley, and Dedon 2014). Our study highlights that APAP promotes a global reprogramming of RNA modifications. The tRNA^Sec^ modification, i^6^A was elevated in WT in response to APAP however this response was again disrupted in the *Alkbh8^Def^* mice. TRIT1 is responsible for modifying position 37 in the anticodon loop of tRNA^Sec^. The i^6^A modification can also be found on various cytoplasmic or mitochondrial tRNA. Cytoplasmic i^6^A modification occurs on tRNA^Ser^ and mitochondrial tRNAs, mtRNA^Cys^ and mtRNA^Ser^ (Schweizer, Bohleber, and Fradejas-Villar 2017). A reduced expression of *Trit1* may account for decreased selenoprotein expression (Warner et al. 2000). Interestingly, an A37G mutation that abolishes the i^6^A37 modification supports a subset of selenoprotein expression including *Txnrd1* and *Gpx4;* however, there was little to no effect on a subset of stress-related selenoproteins such as Gpx1 (Warner et al. 2000). The increase in mcm^5^U and tRNA^Sec^ related modifications suggest that the WT livers respond to APAP by optimizing the production of selenoproteins.

A comparison of saline derived RNA modification data between WT and *Alkbh8^Def^*, revealed that the tRNA modification m^1^A was found at similar levels. After APAP exposure, WT responds by increasing m^1^A modification levels and this reprogramming is disrupted in the *Alkbh8^Def^* mice. m^1^A is present in both cytosolic tRNA species at positions 9, 14, 22,5 7 and 58, and notably on tRNA^Sec^ at position 58, as well as mitochondrial tRNA species at positions 9 and 58 (Ali, Idaghdour, and Hodgkinson 2020; Safra et al. 2017). The addition of the m^1^A modification is catalyzed by the catalytic protein Trmt61A and the RNA binding protein Trmt6 (M. Wang et al. 2016) while the eraser enzyme nucleic acid dioxygenase (Alkbh1) demethylates the m^1^A modification reverting to its unmodified form. Alkbh1 impacts tRNA stability and regulates cleavage dependent on the particular stress response (Rashad et al. 2020). The m^1^A modification aids in mitochondrial tRNA folding and ensures correct structure formation. The m^1^A58 tRNAs have a differential affinity for the elongation factor EF1A which delivers the tRNA into the A-site of the ribosome in protein synthesis (Pan 2018). Our data supports the idea that m^1^A plays an important translational role in protein synthesis in response to APAP and the deficiency of Alkbh8 dysregulates the epitranscriptomic reprogramming. Likely the m^1^A deficiency contributes to the decrease in selenoprotein synthesis, but its decrease could also be globally affecting protein synthesis in the cytoplasm and mitochondria. In addition the m^1^A decrease could be promoting tRNA instability, which could account for the wide-spread decrease in RNA modifications in APAP treated *Alkbh8^Def^* livers.

The m^3^C modification was significantly up-regulated in WT livers after APAP exposure. The m^3^C modification is located at position 32 on various tRNA species as well as their corresponding isoacceptors including; tRNA^Arg^, tRNA^Met^, tRNA^Ser^, tRNA^Thr^ and mt-tRNA^Met^, mt-tRNA^Ser^, mt-tRNA^Thr^ (Cui et al. 2021). The m^3^C modification is catalyzed by the methyltransferase-like protein 6 (METTL6) (Xu et al. 2017) and human METTL6 has been associated with regulating cellular growth, ribosomal occupancy, and pluripotency in hepatocellular carcinoma (HCC). Mettl6 knockout mice displayed altered metabolic activity via altered glucose homeostasis, changes in metabolic turnover, and decreased liver weight suggesting the impact *Mettl6* has on hepatic growth (Ignatova et al. 2020).

### 4.3. Redox and pharmaceuticals responses can be regulated by the epitranscriptome

Oxidative stress is catalyzed by the disruption of redox homeostasis and the subsequent production of ROS. The modifications m^6^A, m^5^C and their respective modifying enzymes have been reported in various cellular models showing the protective role of their dynamic signaling responses. Mettl3, the writer enzyme of m^6^A, has been shown to activate the protective Nrf2 antioxidant pathway in response to colistin-induced oxidative stress in mouse renal tubular epithelial cells (J. Wang et al. 2019). Arsenite induced stress in human keratinocytes resulted in increased expression of other m^6^A writers, WTAP and Mettl14 which regulated changes in gene expression (Zhao et al. 2019) as well as inhibiting p53 activation (Zhao et al. 2020). Eraser and reader enzymes of m^6^A also possess a role in oxidative stress as well. FTO, the eraser enzyme, has implications in mitigating mitochondrial and lipogenesis-induced ROS in HEK293T and clear cell renal carcinoma cells (Zhuang et al. 2019). FTO is a member of the Fe (II)- and α-KG-dependent dioxygenase enzyme family which requires electrons from Fe (II) and α-ketoglutarate to activate a dioxygene molecule (Fedeles et al. 2015). The dioxygenases use oxygen as an electron acceptor to reduce methylation and to form formaldehyde and hydrogen peroxide which may suppress the overproduction of oxidative stress (Chervona and Costa 2012). It is suggested that oxidative stress oxidizes Fe (II) to Fe (III) inhibiting the demethylase activity of eraser enzymes such as FTO (Niu et al. 2015; Ponnaluri, Maciejewski, and Mukherji 2013). However, overexpression of FTO in hepatocytes and myotubes has been linked to increasing lipogenesis and mitochondrial dysfunction leading to increased oxidative stress (Bravard et al. 2011; Guo et al. 2013). Reader enzymes YTHDF1-3 studies have also been shown to have various roles in recognizing oxidative stress signaling and promote regulation of antioxidant pathways, in addition to influencing mRNA stability and translation (Anders et al. 2018; Shi et al. 2019). m^5^C writer enzymes have been shown in HeLa and colon cancer cell lines to upregulate NRF2 and induce cellular senescence upon oxidative stress exposure (Q. Li et al. 2018; Villeneuve et al. 2009). Together with our current Alkbh8 study, past studies highlight the importance of epitranscriptomic marks in response to stress and link mcm^5^U, m^1^A, m^3^C, m^6^A and m^5^C key to the response to ROS.

## Conclusion

Our study highlights that APAP promotes global reprogramming of the epitranscriptome and its effects can be in part modulated by a single epitranscriptome writer (**Fig. 7**). The deficiency of *Alkbh8* resulted in increased liver damage and oxidative stress in *Alkbh8^Def^* mice upon APAP exposure, which can be linked to a global disruption of epitranscriptomic reprogramming, a specific decrease in wobble U modification and the decreased translation of stress response proteins. Chronic exposure of APAP reveals further dysregulation of these critical epitranscriptomic reprogramming efforts, together with global mRNA changes, likely resulting from mitochondrial stress, which leads to additional DNA damage on top of the other pathological consequences. A major finding of our study is that pharmaceuticals promote changes to the epitranscriptome and RNA modification enzymes can modulate outcomes. The efficacy of chemotherapeutics and immunotherapies have also been shown to be modulated by epitranscriptomic systems. METTL3 expression increases resistance to the chemotherapeutic agents 5-fluorouracil, gemcitabine, and cisplatin in pancreatic cancer, through activation of the MAPK signaling pathway (Taketo et al. 2018). The eraser enzyme, FTO, promotes melanoma tumorigenesis and resistance to immunotherapy, while decreased FTO expression increased IFN*y*-induced tumor cell killing and promoted beneficial PD-1 immunotherapy (Yang et al. 2019). Similarly, decreased expression of the eraser ALKBH5 increased the efficiency of PD-1 therapy in both melanoma and colorectal cancer models by decreasing immunosuppressive cells (N. Li et al. 2020). Additionally, wobble U34 modifications catalyzed by ELP3/CTU1/2 have been shown to drive melanomas that express mutated BRAF, with subsequent resistance to anti-BRAF therapy caused by U34-dependent codon-biased translation of metabolic proteins (Rapino et al. 2017). It is important to note that many of the previously published studies showing that the efficacy of therapeutics can be modulated by epitranscriptomic marks were done using *in situ* models and focus on agents specific to cancer treatment. Our study presents an *in vivo* model of how a writer can modulate pharmaceutical stress resulting from a pain reliever and suggests that epitranscriptomic modulation could augment other therapeutics. Overexpression of *Alkbh8* or increased wobble modifications in tRNA^Sec^ could limit APAP toxicity and provide an increased protective role against oxidative stress. Notably, selenium supplementation has been shown to promote increased wobble modification of tRNA^Sec^ and promote selenoprotein expression (Chittum et al. 1997), and it could offer an effective epitranscriptomic enhancement strategy to counteract APAP overdose.

## Supplemental Tables

**Table S1. Measured tRNA modifications in mouse liver tissue after daily dose of APAP (4 Days).** Calculations for each epitranscriptomic mark and comparisons between WT and *Alkbh8^Def^* liver tissue post daily 4 Day exposure to 600 mg/kg of APAP.

**Table S2. WES raw data for all proteins analyzed in 6 hour APAP exposure experiment.** Protein quantitation data was normalized to housekeeping protein, GAPDH, and normalized corrected area analysis setting was set to 100 on ProteinSimple Compass Software.

**Table S3. WES raw data for all proteins analyzed in 4 day APAP exposure experiment.** Protein quantitation data was normalized to housekeeping protein, GAPDH, and normalized corrected area analysis setting was set to 100 on ProteinSimple Compass Software.

**Table S4. All measured epitranscriptomic marks after daily 600 mg/kg APAP for 4 days.** Calculations for each epitranscriptomic gene count and comparisons between WT and *Alkbh8^Def^* liver tissue post 4 day exposure to 600 mg/kg of APAP with reported statistical significance of biological replicates (N = 3) measured by an unpaired t-test. Increased expression (> 0) is reported in red shade and gene counts expressed (< 0) were reported shaded green. Significant comparisons (p ≤ 0.05) are shaded in yellow.

## Supplemental Figures

**Figure S1. Transcripts regulated in *Alkbh8^Def^* Saline vs. WT Saline mice in the 6 hour APAP experiment. (A)** Enhanced volcano plots were generated for *Alkbh8^Def^* saline verse WT saline and **(B)** Metascape analysis were performed for log_2_FC > 2.0 and transcripts regulated were identified.

**Figure S2. Transcripts regulated in *Alkbh8^Def^* Saline verse WT Saline mice in the 4 day APAP experiment. (A)** Enhanced volcano plots were generated for *Alkbh8^Def^* saline verse WT saline and **(B)** Metascape analysis were performed for log_2_FC > 2.0 and transcripts regulated were identified.

**Figure S3. Transcripts regulated in *Alkbh8^Def^* APAP verse WT APAP in the 4 day experiment.** Wildtype and *Alkbh8^Def^* mice (N = 3) were exposed to daily doses of 600 mg/kg APAP over 4 days and tissue was harvested 24 hours after the fourth dose was administered and RNA was purified for analysis by mRNA-seq. **(A)** Enhanced volcano plots were generated for Alkbh8^Def^ APAP verse WT APAP and **(B)** Metascape analysis were performed for log_2_FC > 2.0 and transcripts regulated were identified.

**Figure S4. Selenoproteins measured after 6 hour APAP exposure.** Gpx4, TrxR1 and SelS protein levels in the liver were evaluated using the ProteinSimple WES system. Statistical significance of biological replicates (N = 3) was measured by an unpaired t-test with (*p < 0.05, **p < 0.01, *** p < 0.001).

**Figure S5. Remainder of measured epitranscriptomic marks in mouse liver tissue after APAP (6 hours).** WT and *Alkbh8^Def^* mice (N = 3) were exposed to a single 600 mg/kg dose of APAP and livers were harvested 6 hours after dosing. Modifications were measured using LC-MS/MS. Statistical significance of biological replicates (N = 3) was measured by an unpaired t-test with (*p < 0.05, **p < 0.01, *** p < 0.001).

**Figure S6. Selenoprotein S expression elevated in *Alkbh8^Def^* liver tissue after 4 day APAP exposure.** SelS expression was evaluated using the ProteinSimple WES system. Statistical significance of biological replicates (N = 3) was determined using an unpaired t-test with (*p < 0.05, **p < 0.01, *** p < 0.001).

**Figure S7. tRNA modifications measured after 4 day APAP exposure. (A)** WT and *Alkbh8^Def^* mice (N=3) were exposed to a daily 600 mg/kg dose of APAP and livers were harvested 4 days after dosing. **(B)** Individual bar graphs showing average concentrations of various tRNA chemical modifications measured using LC-MS/MS. Statistical significance of biological replicates (N = 3) were measured by an unpaired t-test with (*p < 0.05, **p < 0.01, *** p < 0.001).

## Acknowledgments

We thank Dr. Sridar Chittur and Marcy Kuentzel from the Center for Functional Genomics at the University at Albany for their help and collaboration with our RNASeq experiments. We also thank Dr. Kevin O’Keefe for his expertise with mouse work along with Ed Zandro M. Taroc and his help with bioinformatic analysis. We finally thank members of the University at Albany RNA Institute and the Nanobioscience constellation at SUNY Poly’s College of Nanoscale College and Engineering for their insightful discussion and feedback.

