## Supplemental Figures for "Epitranscriptomic reprogramming is required to prevent stress and damage from acetaminophen"

#### Supplemental Figure S1.

A)

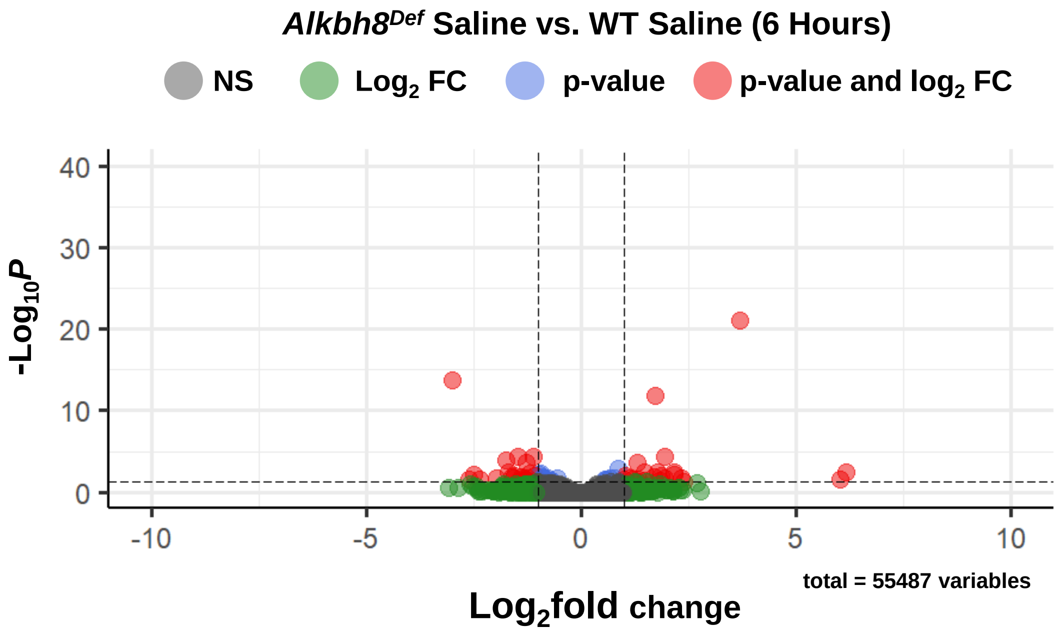


B)

### Up-regulated

| Ensembl ID | Gene Symbol | Description | Biological Process (GO) |
|---|---|---|---|
| ENSMUSG00000038754 | **Elovl3** | elongation of very long chain fatty acids (FEN1/Elo2, SUR4/Elo3, yeast)-like 3 | GO:0019367 fatty acid elongation, saturated fatty acid;GO:0019368 fatty acid elongation, unsaturated fatty acid;GO:0034625 fatty acid elongation, monounsaturated fatty acid |
| ENSMUSG00000025229 | **Pitx3** | paired-like homeodomain transcription factor 3 | GO:1904935 positive regulation of cell proliferation in midbrain;GO:1904933 regulation of cell proliferation in midbrain;GO:0033278 cell proliferation in midbrain |
| ENSMUSG00000037583 | **Nr0b2** | nuclear receptor subfamily 0, group B, member 2 | GO:0032922 circadian regulation of gene expression;GO:0032024 positive regulation of insulin secretion;GO:0090277 positive regulation of peptide hormone secretion |
| ENSMUSG00000044646 | **Zbtb7c** | zinc finger and BTB domain containing 7C | GO:1903025 regulation of RNA polymerase II regulatory region sequence-specific DNA binding;GO:2000677 regulation of transcription regulatory region DNA binding;GO:0045600 positive regulation of fat cell differentiation |
| ENSMUSG00000026077 | **Npas2** | neuronal PAS domain protein 2 | GO:1903367 positive regulation of fear response;GO:2000987 positive regulation of behavioral fear response;GO:0051775 response to redox state |
| ENSMUSG00000038508 | **Gdf15** | growth differentiation factor 15 | GO:0060400 negative regulation of growth hormone receptor signaling pathway;GO:0060398 regulation of growth hormone receptor signaling pathway;GO:0002023 reduction of food intake in response to dietary excess |

### Down-regulated

| Ensembl ID | Gene Symbol | Description | Biological Process (GO) |
|---|---|---|---|
| ENSMUSG00000066687 | **Zbtb16** | zinc finger and BTB domain containing 16 | GO:0048133 male germ-line stem cell asymmetric division;GO:0051138 positive regulation of NK T cell differentiation;GO:0042078 germ-line stem cell division |
| ENSMUSG00000048794 | **Cfap100** | cilia and flagella associated protein 100 | GO:0008150 biological_process |
| ENSMUSG00000091898 | **Tnnc1** | troponin C, cardiac/slow skeletal | GO:0032972 regulation of muscle filament sliding speed;GO:0002086 diaphragm contraction;GO:0003011 involuntary skeletal muscle contraction |
| ENSMUSG00000024365 | **Cyp21a1** | cytochrome P450, family 21, subfamily a, polypeptide 1 | GO:0006705 mineralocorticoid biosynthetic process;GO:0008212 mineralocorticoid metabolic process;GO:0006704 glucocorticoid biosynthetic process |

#### Supplemental Figure S2.

A)

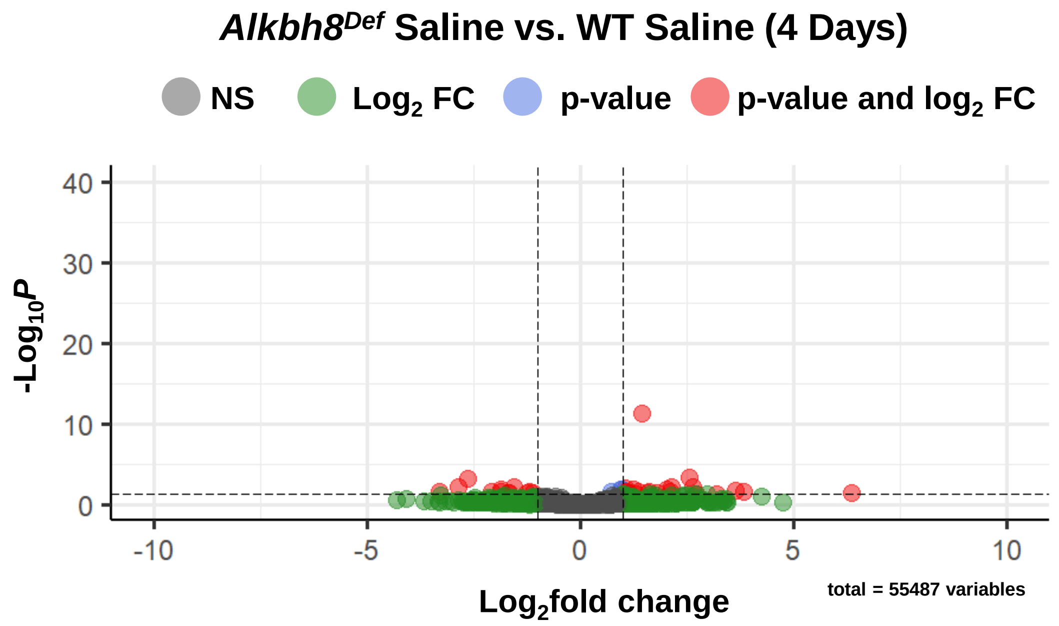


B)

##### Up-regulated

| Ensembl ID | Gene Symbol | Description | Biological Process (GO) |
|---|---|---|---|
| ENSMUSG00000035498 | **Cdcp1** | CUB domain containing protein 1 | GO:0008150 biological_process |
| ENSMUSG00000022351 | **Sqle** | squalene epoxidase | GO:0140042 lipid droplet formation;GO:0034389 lipid droplet organization;GO:0016126 sterol biosynthetic process |
| ENSMUSG00000041782 | **Lad1** | ladinin | |
| ENSMUSG00000025229 | **Pitx3** | paired-like homeodomain transcription factor 3 | GO:1904935 positive regulation of cell proliferation in midbrain;GO:1904933 regulation of cell proliferation in midbrain;GO:0033278 cell proliferation in midbrain |
| ENSMUSG00000024036 | **Slc37a1** | solute carrier family 37 (glycerol-3-phosphate transporter), member 1 | GO:0015712 hexose phosphate transport;GO:0015760 glucose-6-phosphate transport;GO:0035435 phosphate ion transmembrane transport |
| ENSMUSG00000026077 | **Npas2** | neuronal PAS domain protein 2 | GO:1903367 positive regulation of fear response;GO:2000987 positive regulation of behavioral fear response;GO:0051775 response to redox state |
| ENSMUSG00000025185 | **Loxl4** | lysyl oxidase-like 4 | GO:0018057 peptidyl-lysine oxidation;GO:0018158 protein oxidation;GO:0030199 collagen fibril organization |
| ENSMUSG00000023067 | **Cdkn1a** | cyclin-dependent kinase inhibitor 1A (P21) | GO:1905178 regulation of cardiac muscle tissue regeneration;GO:1905179 negative regulation of cardiac muscle tissue regeneration;GO:0061026 cardiac muscle tissue regeneration |

##### Down-regulated

| Ensembl ID | Gene Symbol | Description | Biological Process (GO) |
|---|---|---|---|
| ENSMUSG00000059060 | **Rad51b** | RAD51 paralog B | GO:0010971 positive regulation of G2/M transition of mitotic cell cycle;GO:1902751 positive regulation of cell cycle G2/M phase transition;GO:0001832 blastocyst growth |
| ENSMUSG00000038060 | **Dlec1** | deleted in lung and esophageal cancer 1 | GO:0008285 negative regulation of cell proliferation;GO:0042127 regulation of cell proliferation;GO:0008283 cell proliferation |
| ENSMUSG00000022528 | **Hes1** | hes family bHLH transcription factor 1 | GO:0061105 regulation of stomach neuroendocrine cell differentiation;GO:0061106 negative regulation of stomach neuroendocrine cell differentiation;GO:1905933 regulation of cell fate determination |

**Supplemental Figure S3.**

A)

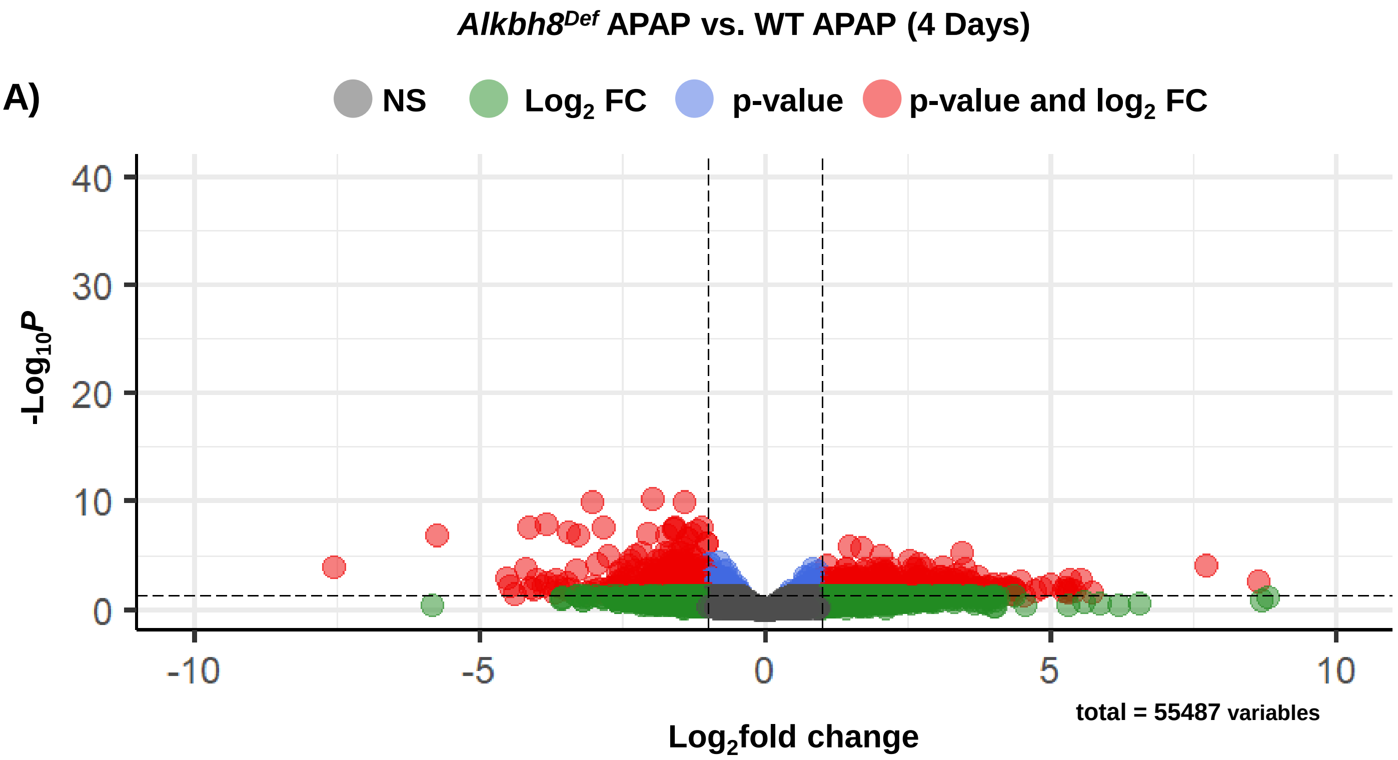


B)

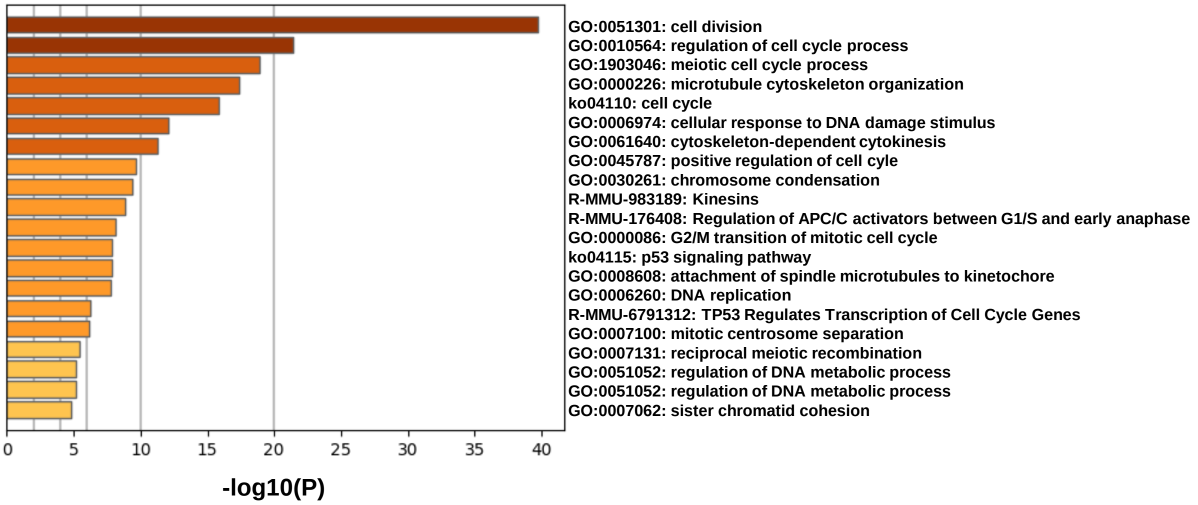

#### Supplemental Figure S4.

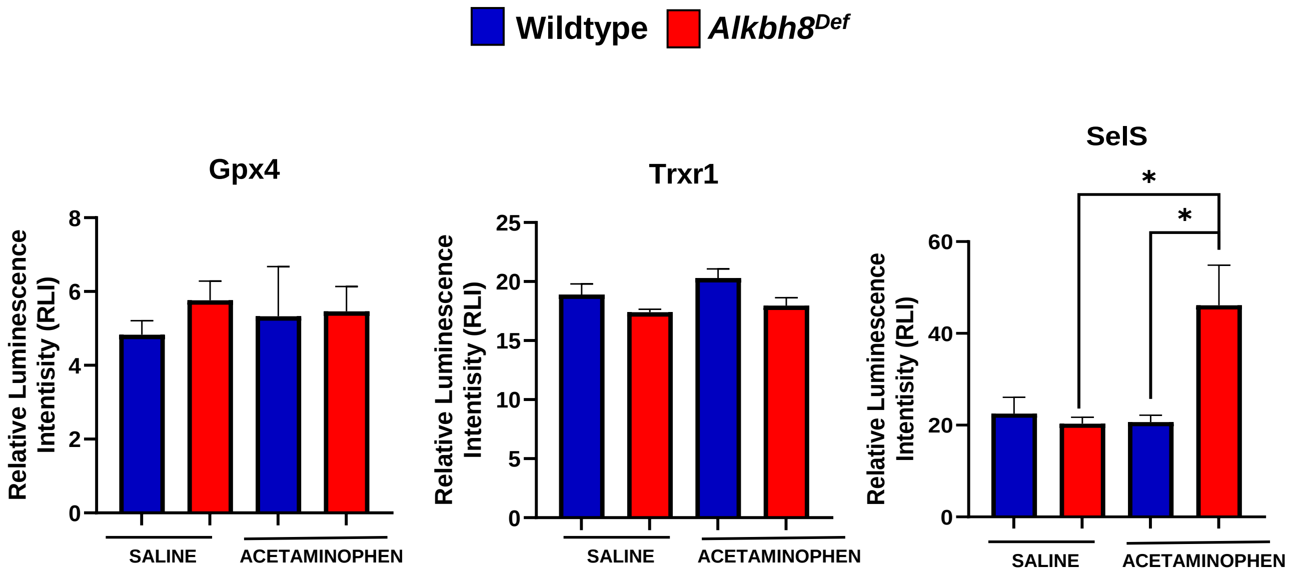

### Supplemental Figure S5.

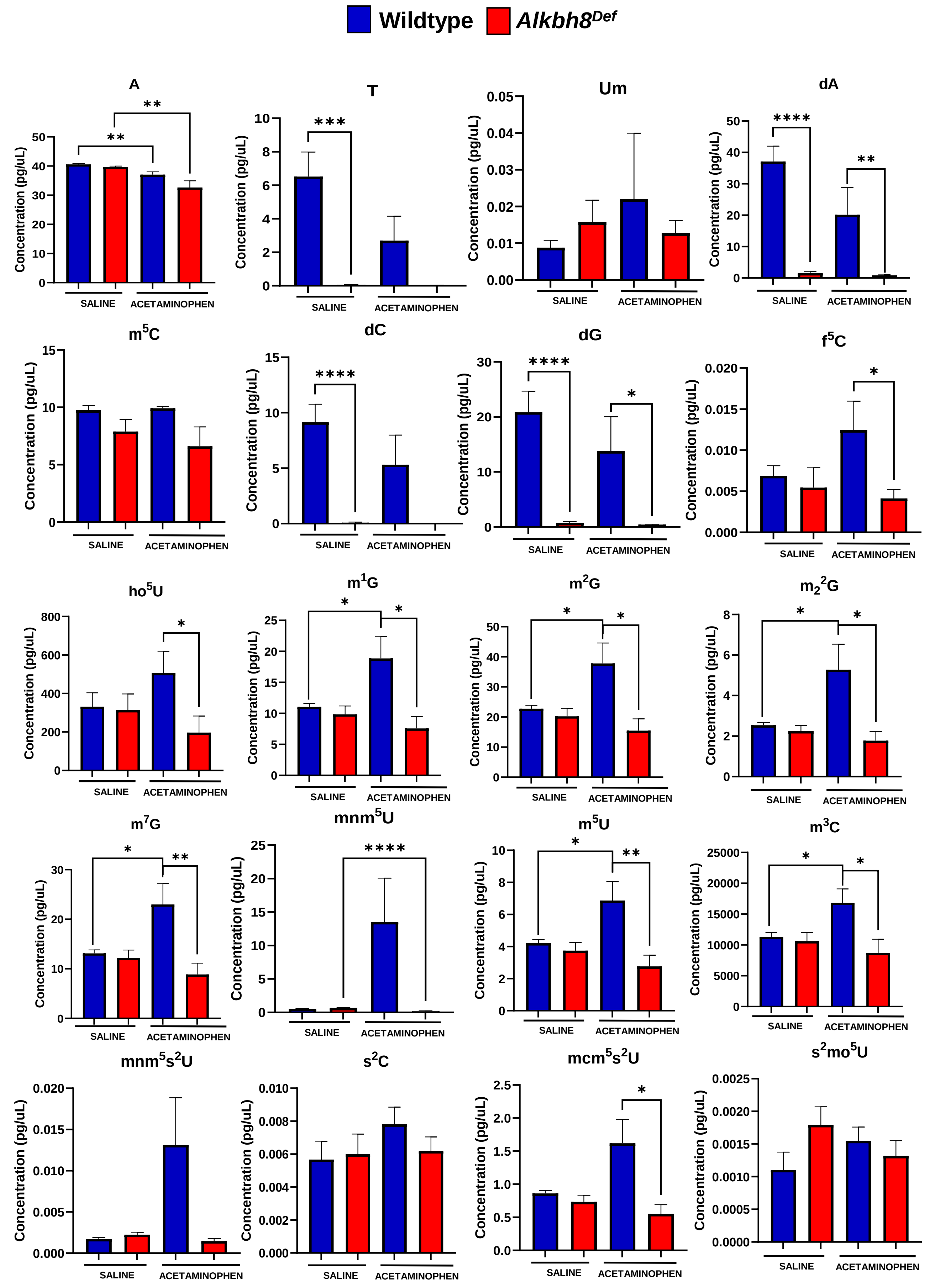

### Supplemental Figure S6.

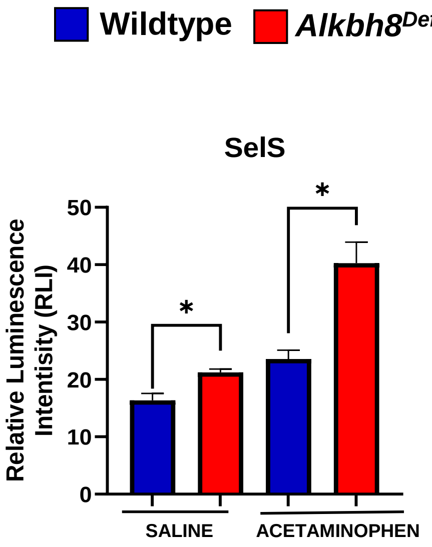

### Supplemental Figure S7.

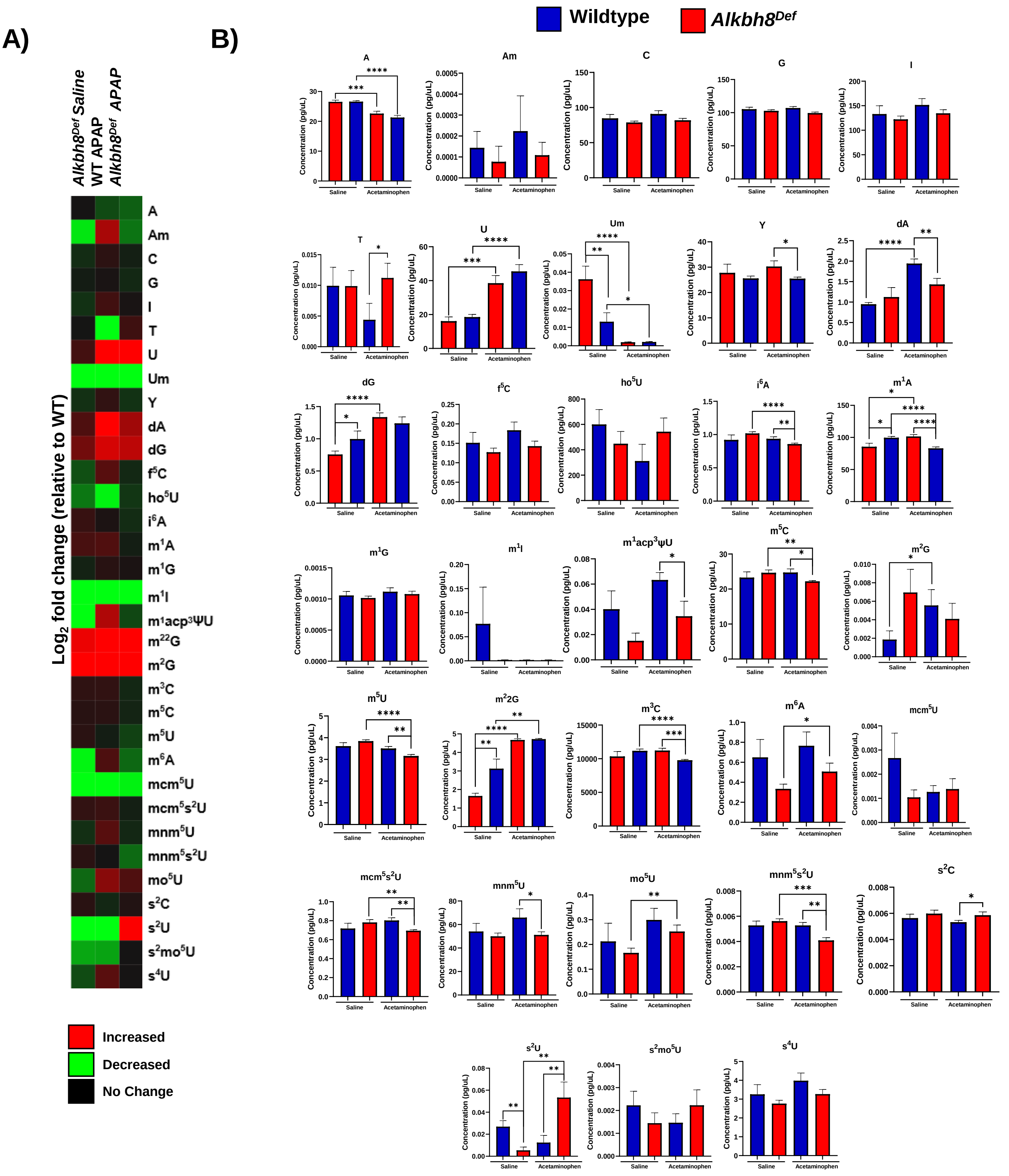
